# The reconstruction and functional mapping of a recurrent microcircuit in *Drosophila* mushroom body

**DOI:** 10.1101/819227

**Authors:** Guangxia Wang, Bangyu Zhou, Shengxiong Wang, Kai Yang, Jianjian Zhao, Xing Yang, Yiming Li, Lijun Shen

**Affiliations:** Institute of Neuroscience and State Key Laboratory of Neuroscience, Center for Excellence in Brain Science and Intelligence Technology, Chinese Academy of Science, Shanghai 20003, China; University of Chinese Academy of Sciences, Shanghai 200031, China; State key Laboratory of Brain and Cognitive Science, Institute of Biophysics, Chinese Academy of Science, Beijing 10010, China; Institute of Automation, Chinese Academy of Sciences, Beijing 10090, China

## Abstract

In *Drosophila melanogaster*, mushroom body and anterior paired lateral (APL) neurons play important roles not only in learning and memory but also in high cognitive behavior, reversal learning. The circuit between APL neurons and Kenyon cells (KCs) in the mushroom body underlies this behavior, including reversal learning, and electron microscopy (EM) methods must be used to reveal this circuit. Here, we reconstructed the connections between mushroom body cells and APL neurons in the vertical lobe of the mushroom body via focused ion beam scanning electron microscopy (FIB-SEM) and sparse genetic horseradish peroxidase (HRP) labeling. We offer the first EM evidence that recurrent network and lateral inhibition connections exist between APL neurons and KCs in the vertical lobe of the mushroom body. This circuit is the neural basis of action selection decision making, associative learning and reversal learning. Additionally, dopamine neurons project to different areas of mushroom bodies and, together with extrinsic neurons and KC axons, form a compartmental structure of mushroom body axons, thereby restricting the KC-mushroom body output neuron (MBON) response to local compartments. Whether APL neurons also respond locally is uncertain. We found that APL neurons exhibited input and output synapses that were intermixed and arranged on enlarged and thin sections, respectively, resembling a string of beads. Different KCs were found to project to APL neurons nonrepetitively, forming a local circuit structure. Furthermore, using a single neurite calcium imaging method, we identified local calcium domains on this circuit, suggestive of individual electrical compartments. The electrically recorded APL neurons were nonspike neurons that selectively responded to odor in both the lobes and calyx. Thus, the localized APL neuron responses coordinate with mushroom body–dopamine-MBON compartmental function.

## Introduction

Mushroom bodies are the most important structures in *Drosophila* and are primarily composed of KCs (KCs). The cell bodies and dendrites of three different intrinsic neurons constitute the calyx, and the axons constitute the peduncle and three lobes, the αβ, α′β′, and γ lobes (Tanaka, Tanimoto, & Ito, 2008). Mushroom bodies are the third level of the olfactory system; they receive input from projection neurons in the calyx and send information through contacts with extrinsic neurons in the lobes (Aso et al., 2014a, 2014b).

A pair of symmetrically distributed GABAergic interneurons, called anterior posterior lateral (APL) neurons, each of which sends out two branches, innervate both the calyx and lobes of mushroom bodies. APL neurons are thought to function similarly to their homolog giant GABAergic neurons (GGNs) in the locust, which receive information in the calyx of the mushroom body and send global feedback inhibition to KCs in the lobes (Papadopoulou, Cassenaer, Nowotny, & Laurent, 2011). APL neurons and mushroom bodies participate in sparse odor coding (Lei, Chen, Li, Liu, & Guo, 2013), olfactory learning inhibition (Liu & Davis, 2009), mutual odor inhibition, visual reversal learning (Ren, Li, Wu, Ren, & Guo, 2012), olfactory reversal learning (Wu, Ren, Li, & Guo, 2012), decorrelated odor coding (Lin, Bygrave, de Calignon, Lee, & Miesenbock, 2014), and visual decision making. Both blocking APL neuron transmission and decreasing the expression of GABAA receptors in the αβ lobe of mushroom bodies inhibit visual reversal learning in *Drosophila* (Ren et al., 2012). Recent studies have demonstrated that APL neurons coexpress octopamine and GABA (Pitman et al., 2011). Further studies found that octopaminergic APL neurons project to the α′β′ lobe of mushroom bodies in the mushroom body output neuron (MBON)-β′2mp pathway and serotonergic dorsal paired medial (DPM) neurons project to the αβ lobe of mushroom bodies in the MBON-β2β′2a pathway to mediate 3-h anesthesia-sensitive memory (Pitman et al., 2011; Yang et al., 2016).

Some neuromodulator neurons innervate mushroom bodies, and dopaminergic neurons might mediate unconditional stimuli, such as reward and punishment during learning. Dopamine neurons couple with KCs, which encode odor information for conditioning, learning and storage of olfactory information.

Dopaminergic and extrinsic neurons have been shown to project to different areas of mushroom body lobes and separate mushroom bodies into 15 functional compartments (Aso et al., 2014a, 2014b). The dopamine neurons that project locally encode innate states as well as external experiences. Together with KC axons and extrinsic neurons, the locally projecting dopamine neurons form local information circuits in this area (Cohn, Morantte, & Ruta, 2015), dopamine neurons constrain KC neuron responding locally.

Studies on APL neuron morphology have shown that these neurons innervate whole lobes and the calyx (Liu & Davis, 2009), and both presynapses and postsynapses of APL neurons are found in all these areas. APL neurons with KC neurons play roles in numerous high cognitive behaviors, and the following questions remain unanswered: What are the neural circuits of APL neurons and KCs that underlie these behaviors? Does a local circuit also exist between APL and KCs? Since DA-KC-MBON circuit function locally, APL neurons intermixed with mushroom body are also well segmented via DA neurons. Moreover, APL neurons are activated via KC neurons which have locally response at the vertical lobe. Do APL neurons function locally at lobe and calyx?

To explore these questions, we introduced a 3D electron microscopy (EM) reconstruction method to obtain information at the nanoscale. We genetically labeled APL neurons with horse-radish peroxidase (HRP) and examined the circuit using focused ion beam series electron microscopy (FIB-SEM). We obtained substantial amounts of continuous EM data in the mushroom body vertical lobe (volume EM data) and then reconstructed APL neurons and their connectomes. We found that APL neurites received KC input and then KC sent out information to APL neurites, forming a recurrent loop. We showed that individual APL neurites have intermixed presynapses and postsynapses that are found on alternate arranged enlarged and thin segments, respectively. Thus structure constitutes the electrically separation on APL single neurite, forming a local information flow from KC to enlargement of APL and send out information from thin part of APL to other KC. The structural evidence supports the potential existence of a local circuit between APL neurons and KCs. Electrophysiological recording revealed that APL neurons elicit nonspikes. In addition, via a calcium imaging method used to indicate neuron activity, we observed calcium domains on individual APL neurites, suggesting that signal propagation is limited on these neurites and that this electrical segmentation is underscored by the enlargement and thinning of the neurite. APL neurons send out odor-dependent signals locally in both the calyx and the lobes. These data support the hypothesis that a local circuit but not a global circuit exists between APL neurons and KCs in mushroom bodies, which complements the compartmental structure of mushroom bodies and might enable the various inhibitory local circuits associated with learning and innate behavior and other behavior. Thus, we supply the first EM evidence that local APL-KC neurons forms a recurrent network that is the neural basis of learning, reversal learning, and decision-making behavior.

## Results

### Three-dimensional reconstruction of HRP-labeled APL neurons and their connections in the mushroom body vertical lobe

To explore the specific APL neurons and their connectomes, volume EM reconstruction was applied. Because several high cognitive behaviors are based in the vertical lobe, we explored the connectomes in this region. Two strategies are available for studying a target neuron and its connections: one strategy is constructing all of the neurons in a specific brain area and then identifying the target neuron and its connections (Macpherson et al., 2015), thus dense reconstruction. The other strategy is sparse reconstruction (Kasthuri et al., 2015), in which neurons of interest are sparse labeled and their connections are identified directly. The second strategy is much less time consuming, providing us with the opportunity to identify our circuit of interest. In our study, the classic EM tracer HRP was applied as a labeling method. We drive HRP (*UAS-HRP-mCD8-GFP*) expression via *GH146-Gal4* to specifically labeling APL neurons at the mushroom body. HRP catalyzes the formation of a 3,3’-diaminobenzidine (DAB) polymer (Joesch et al., 2016; Kasthuri et al., 2015), which binds osmium to increase the electron density. FIB-SEM then enabled us to obtain volume EM data of the vertical lobe of mushroom body from tip to bottom, approximately 30 (X) × 37 (Y) × 52 (Z) µm^3^ (Fig. S1) and 40 nm thick per image.

The APL neurons expressing HRP exhibited deeper and darker membranes than other neurons after EM preparation (Fig. 1A1 and A2), and our serial EM data allowed for synapse identification across several layers (Fig. 1B1-B6). With a work group comprising 5 people, we applied strict standards to identify synapses and neurons across layers. Drosophila synapses contains three components: a T-bar and vesicle at the presynpase and a darkened membrane at the postsynapse (Marta et al., 2011), which have been identified across several layers of EM data. Some labeled APL neurons showed darkened membranes that continued across several layers of the EM data, whereas others showed darkened membranes that reappeared intermittently across layers. We marked the above as the APL neurons in our reconstruction work as criteria. To ensure that all data identified were accurate, we built a semiautomated workflow (Fig. S2, *Supplemental Material*). Only 100% positive data (all 5 people confirmed the data) were saved. After reconstructing an APL neuron and its pre- and postsynapses, we continued reconstructing the presynapse and postsynapse neuron skeleton of the APL neuron with the same standards. In the end, we obtained 3D reconstruction data for an APL neuron and its connection neurons in the vertical lobe (Fig. S3, Fig. 1G). In total, all reconstructed neurons and connections totaled 72432 nodes (nodes are labeled as the skeleton of neurons on each layer of EM data), 3968 µm of neurites, 183 neurites and 601 synapses (Table 1).

**Fig. 1.**
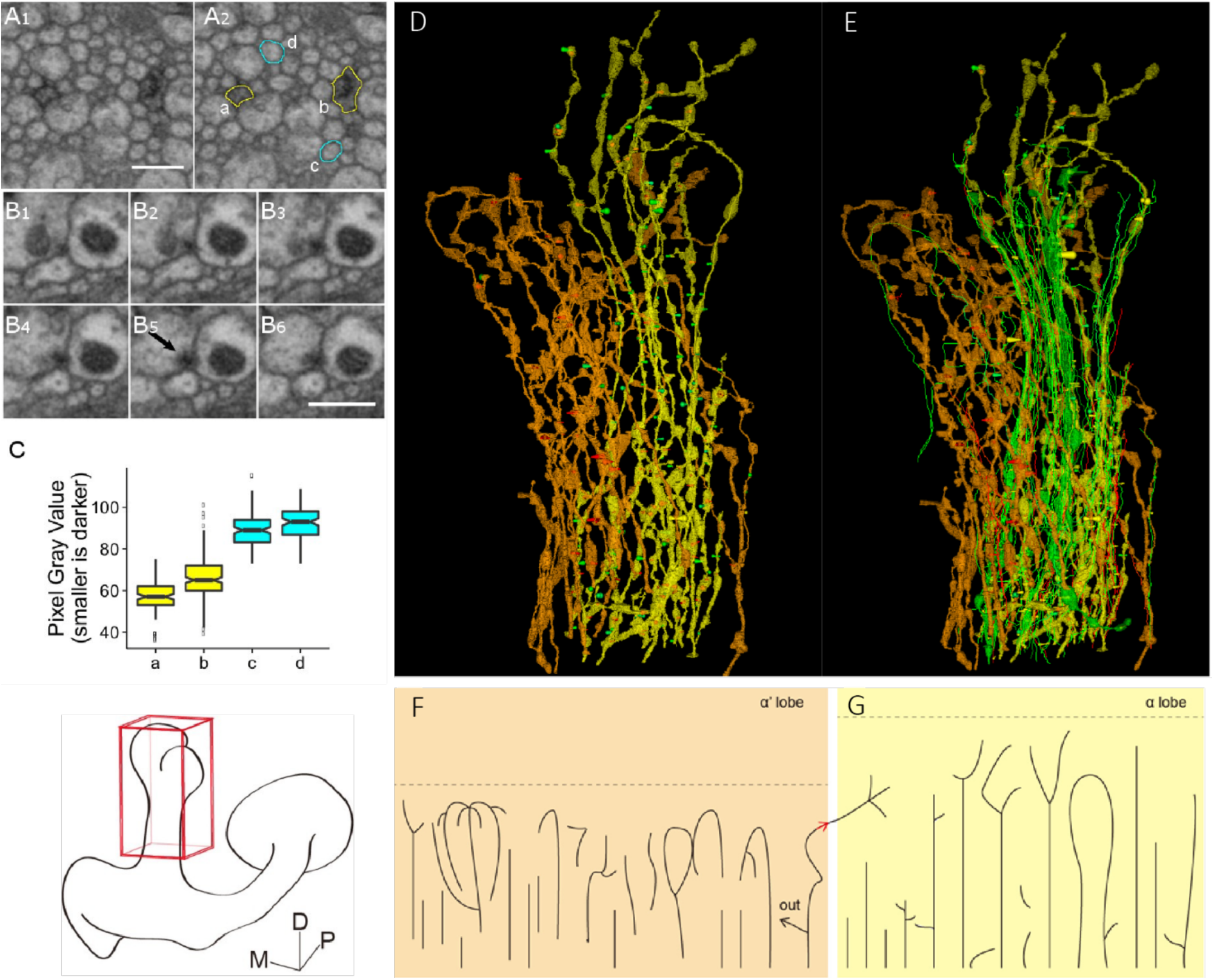
Reconstruction of genetically labeled APL neurons and their connections in the mushroom body vertical lobe. **(A)** Two APL neurite branches with darkened membranes. In A2, yellow circles indicate membranes labeled with HRP in A1, while cyan circles indicate those not labeled. **(B)** Continuous synapse with T-bars. (B5) The arrow indicates a typical T-bar on this layer. The scale bar is 50 nm. **(C)** Pixel gray value measured in areas a, b, c, and d in A2. a and b are both darker than c and d. The scale bar is 500 nm. **(D)** 3D reconstructed APL neurite. **(E)** 3D reconstructed APL neurite and its connected neurite. **(D)** Top, dorsal. An APL neurite in the α lobe is shown in yellow, and an APL neurite in the α′ lobe is shown in orange. APL neurites in the α and α′ lobes have different trends, and they are sparse at the top of the mushroom body vertical lobe and dense in the middle. The green cone represents APL neurite input, and the red cone represents APL neurite output. **(E)** Those overlapping areas are not connected. In addition, the green neurite represents input neurons of APL neurites, and the red neurite represents output neurons of APL neurites. A few bidirectional neurons are also labeled green. Only skeletons of those neurons are displayed. **(F)** APL neurites in the α′ lobe. All constructed data contain 15 independent neurites longer than 12 µm. At the tip of the neurite, there are some horizontal connections and some complex branches, including a circle branch. There is one branch that receives input from neurons out of the mushroom body (labeled by the black arrow and the word out) and also projects to the α lobe. **(G)** APL neurites in the α′ lobe. There are 11 independent branches longer than 12 µm. Most APL neurites are vertical with simple branches, and the vertical branches have few horizontal connections unless near the top of the mushroom body vertical lobe. APL neurites less than 4 µm are omitted from this figure but had no branches.

**Table 1.** Reconstruction data of APL connections in the mushroom body vertical lobe. (*) nodes represent the annotation of the APL skeleton, which is the component of the skeleton on each layer. (**) represent the reconstruction results; not all synapses have an output neurite.

| MB lobe | Neuron | Length ($\mu\text{m}$) | Nodes* | Neurites | Input synapse** | Output synapse** |
| --- | --- | --- | --- | --- | --- | --- |
| $\alpha$ | APL | 680 | 12190 | 22 | 96 | 103 |
|  | Input | 1750 | 34090 | 71 | 60 | 198 |
|  | Output | 332 | 6918 | 38 | 44 | 41 |
|  | Bidirectional | 129 | 2646 | 4 | 7 | 17 |
| $\alpha'$ | APL | 930 | 13984 | 30 | 14 | 27 |
|  | Input | 72 | 1273 | 8 | 0 | 8 |
|  | Output | 75 | 1331 | 10 | 10 | 0 |

APL neurites in the α and α′ lobes are independently, separated and exhibited obviously different trends (Fig. 1D). APL neurons penetrated into the inside of the vertical lobe of the mushroom body and dispersed more evenly in the horizontal direction. Overall, APL neurites in the vertical lobe do not appear as a complete net structure at we thought in previous study, at least in the α lobe. Firstly, we found only one connection between APL neurites in the α and α′ lobes, which also connected with APL branches going into the vertical lobe of mushroom body, and astrocytes are find to isolate α and α′ lobes under EM structure. Secondly, APL neurons in the top of the data set, i.e., the dorsal side of the vertical lobe of the mushroom body, were more sparse in the vertical direction. Fig. S4). A large proportion of these neurites extended in the vertical direction (Fig. S5), consistent with the direction of the mushroom body neurons around them. APL neurites in the α lobe extended straight upwards, while those in the α′ lobe spiraled around those in the α lobe. Thirdly, there were more branches at the tip of the vertical lobe and no branches at the stalk, especially in the α′ lobe. They are rarely separated and dependent neurites at the stalk of the vertical lobe. These observations suggest that APL neurites in the α′ lobe have much more complex connections than those in the α lobe. APL neurites in the α′ lobe extended more horizontally (Fig. S4), but the radius of APL neurites was approximately 10 nm near the section interval, making tracing the APL horizontal branches in the α′ lobe difficult. Therefore, we obtained less data on APL connections in the α′ lobe. In total, we observed 22 APL neurites in the α lobe, 30 APL neurites in the α′ lobe, 113 neurites connecting with APL neurons in the α lobe, and 18 neurites connecting with APL neurons in the α′ lobe (Table 1). Notably, there were fewer than 20 APL branches in the horizontal section of the α lobe (Fig. S4, Table 1) and approximately 1000 KCs in the α lobe; in contrast, there were fewer than 30 APL branches in the α′ lobe (Fig. S4, Table 1) and approximately 300 KCs in the α′ lobe. These data suggest that the average number of APL branches in the α lobe needs to be at least 50 to cover all KCs, while the number of branches in the α′ lobe needs to be at least 10. The horizontal neurite of APL, less than 40nm, might be missed in this dataset due to it below the lateral resolution of EM. While indeed they are few in reality, because there are fewer neurites larger less than 40nm, so neurites less than 40nm are rare. Evenly they exsits, the ability of chemical signal transduction could be ignored due to the thinnest radius. Such, APL could not send signal globally but locally.

Due to the relative comprehensive reconstruction data of APL and its connections in the α lobe, we further studied their neural connections next. We classified neurons connecting with APL neurites into 3 groups based on the information flow between these neurons and APL neurons. Neurons projecting to APL neurons were called input neurons, those that receive input from APL neurites were called output neurons, and those that both project to APL neurites and receive information from APL neurites were called bidirectional neurons. Among the input and output neurons, over 60% formed a single contact with APL neurons. Those neurons extended along APL neurons vertically in a bunch and then separated from APL neurons after forming a synaptic contact at a distance. A few input neurons also formed two or three synaptic contacts with APL neurons (Table 1).

In our reconstructed results, the total number of input synapses (presynapses) in the α lobe was only approximately 100 (Table 1), which is far fewer than the total number of input synapses needed to cover all KCs; even considering that our results underestimate the true situation by four to five times (Fig. S6), we did not find enough input synapses to cover all KCs. Similar conditions were observed in the output synapses (postsynapses) of APL neurons in the α lobe and α’ lobe. Thus, unlike the GGN in locusts, APL neurons in the vertical lobe do not seem to act as broad inhibitory neurons, broadcasting inhibitory signals to all KCs, suggesting that the physiological characteristics of GGNs may not be directly applicable to those of APL neurons.

### APL pre- and postsynapses correlate with enlarged and thin neurite regions, respectively

The neurites of APL neurons resembled a string of beads (Fig. 2A) with intermixed enlarged and thin regions arranged on a single neurite, just as observed in the calyx (*unpublished data*). Presynapses were mainly located in the enlarged regions, whereas postsynapses were mainly located in the thin regions on APL single neurite (Fig. 2). Furthermore, mitochondria, which supply ATP for synapse release and vesicles, were located at the enlarged sections.

**Fig. 2.**
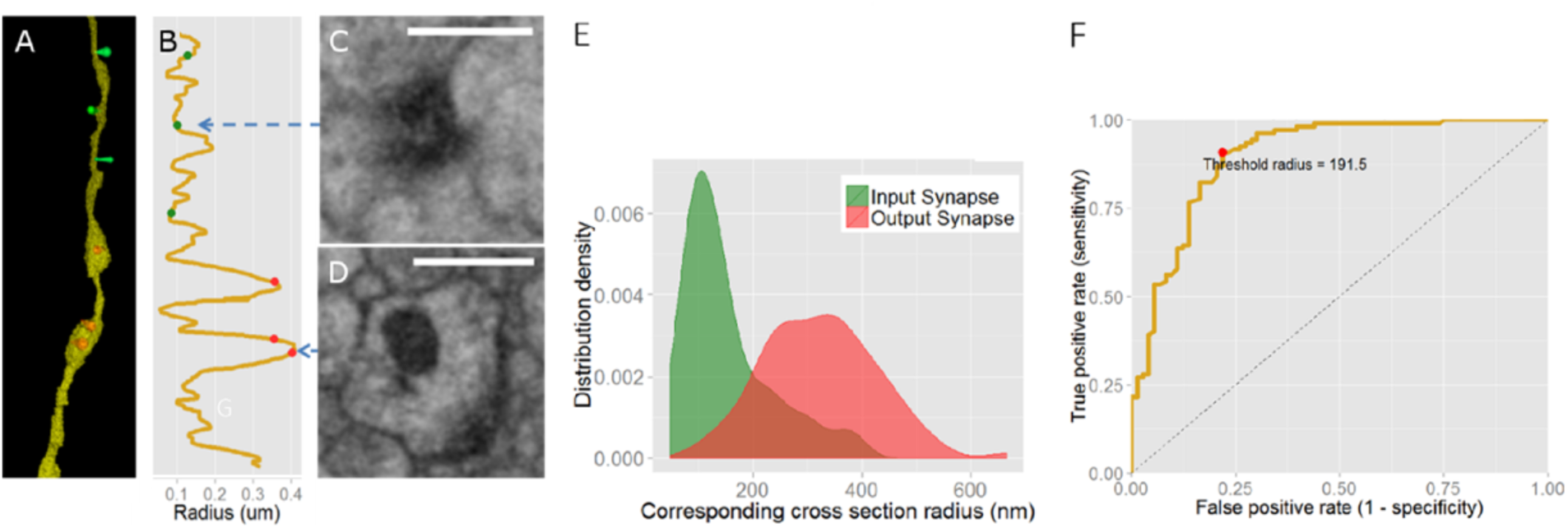
Pre- and postsynapses located at both enlarged and thin regions of APL neurites. **(A)** 3D reconstructed segment of a single APL neurite, overall resembling a string of beads with intermixed enlarged and thin regions. The green cones indicate APL neurite input, while the red cones indicate APL neurite output. **(B)** Illustration of the section radius of the APL neurite in (A). The vertical axis represents the relative location of the APL neurite, and the horizontal axis represents the section radius of the APL neurite perpendicular to the relative location. In both A and B, the green cones represent input synapses, and the red cones represent output synapses. **(C)** A postsynapse at the thin part of the APL neurite. **(D)** A presynapse at the enlarged part of the APL neurite. In (C) and (D), the T-bar structure is distinguishable. The scale bar is 500 nm. **(E)** Distribution densities of section radii with input and output synapses across the APL neurite. The green curve represents the distribution density of section radii at input synapses. Most radii are approximately 100 nm, while a small number are larger. The red curve represents the distribution density of section radii at output synapses. The radii are widely distributed near 200 to 400 nm. **(F)** ROC curve to assess the direction of synapses using neurite section radii as the classifier. The curve shows a steep upward convex shape, indicating a high predictive accuracy of the direction using the synaptic radius. The red dot indicates the best classification threshold, 191.5 nm. The diagonal line represents the performance of random classifiers to determine the direction of each synapse.

We then examined whether the enlarged regions correlate with the location of APL neurite presynapses. To answer this question, we first imaged the presynaptic proteins of APL neurons under a light microscope with *GH146-Gal4* driving syb-GFP expression, revealing sparse puncta of syb-GFP similar to those of the enlarged regions of APL neurons, and it hint a colocalization between presynapse protein and enlargement part of APL neuorns (Fig. S9 and S10). We next investigated the relationship between the radii of the APL neurites and synapse direction using the EM dataset. We analyzed the distribution densities of input synapses (presynapses) and output synapses (postsynapses) across APL neurite radii. In our dataset, the input and output synapse peaks were clearly separated, with input synapses mainly centered at small radii (100 nm) and output synapses mainly centered at large radii (300 nm) (Fig. 2E), indicating a correlation between APL neurite radii and synapse direction. Therefore, we examined whether the radius of an APL neurite could predict the direction of the synapse. To answer this question, we applied receiver operating characteristics (ROC) analysis to assess synapse direction using the section radius of the neurite as the classifier (Fig. 2F). The upward, steep shape of the ROC curve indicated a high predictive accuracy, with the best separation point at 191.5 nm. This result shows that the radii of APL neurites were highly correlated with synapse direction and that synapse direction could be predicted based on neurite radius.

A previous study showed that pre- and postsynapses are located in both the lobes and the calyx, but whether they are located on the same neurite or on different neurites was unclear. Our study confirmed that these synapses are located on the same neurite.

To display the mixed arrangement of the synapses in different directions of APL neurons and compare among neurons, we excluded the distance between synapses and the length and branches of neurons and extracted the topological arrangement between them. Here, we developed an “*arrangement index*” (AI) to achieve this goal (*Supplemental Material*). Compared with A17 neurons in mammalian retina, the synaptic alignment of APL neurons was slightly less mixed (Fig. S7).

Another parameter we applied is the distance between synapses (Fig. S8). For each synapse, the shortest distance to the nearest synapses was identified and recorded. Similarly, the shortest distance between two input synapses was identified in the “input-input” group, the shortest distance between two output synapses was identified in the “output-output” group, and the shortest distance between an input synapse and output synapse was identified in the “input-output” group. Comparing these three sets of data, we found no significant differences in the distribution of the distances, with all being approximately 2 µm. This lack of a difference among these dataset further describes the mixed arrangement of the synapses; if input and output synapses were separated from each other, the distance in the input-output group would be significantly greater than that in the input-input and output-output groups. Therefore, we conclude that the input and output synapses in APL neurons are arranged in a mixed manner.

Importantly, electrical transduction properties differ between thin and enlarged neurites, according to a previous study (Grimes, Zhang, Graydon, Kachar, & Diamond, 2010; Rall, 1969), with relatively thin neurites exhibiting low conductance. Therefore, the alternating thin and enlarged neurite segments result in a separation of the neuronal activity. Thus is the electrically signal separation basis on Amacrine A17 neurons which have broad branches with varicosities and thin neurites alternate arranged on it, and feedback inhibition to presynaptic biopolar cells located at the varicosities. Thus, APL neurites receive input at enlarged sections and provide output from thin sections, thus constructing a basic local neural circuit unit. Furthermore, APL neurons formed reciprocal connections with neighboring neurons (Fig. 2E), but the identity of these connections neurons required further exploration.

### APL neurons mediate the lateral inhibition of KCs in the vertical lobe and the recurrent network between APL and KCs

Together with APL neurons, six subtype of KCs, several dopamine neurons, some MBONs and DPM neurons project to the mushroom body and the vertical lobes. A study of the dopamine 2 receptor, which supports anesthesia-resistant memory, showed that dopamine neurons make synaptic connections with APL neurons via the D2R receptor (Scholz-Kornehl & Schwarzel, 2016). Gap junctions exist between APL neurons and DPM neurons (Pitman et al., 2011; Wu et al., 2011). Blocking the output of KCs completely silences APL neuron activity, suggesting a direct or indirect connection between APL neurons and KCs (Kageyama & Meyer, 1987; Lin et al., 2014). These studies under light microscopy to demonstrate that various neurons are connected to APL neurons, but EM evidence is still lacking. To deconstruct the neural circuit comprising APL and other neurons under EM, the other neurons must first be identified.

These reconstructed neurons under EM exhibit specific morphologies under light microscopy that separate them from each other. KCs have no branches (Lee, Lee, & Luo, 1999), while other neurons have extensive branches in the vertical lobe (Aso et al., 2014a, 2014b). Hence, neurons connected to APL neurons were classified based on their proportion in vertical lobe and morphology. Morphology of APL neurons mainly consider two features, the length of neurites and the number of bifurcations (Fig. 3CD). Neurites belong to KCs if they have no branches in the vertical lobe and are of a length equal to or slightly less than that of the total vertical lobe. The priori probability of KC for these reconstructed neurons is very high, considering that 1300 KC neurons account for more than 90% of them in the vertical leaves. Neurites that have more than one branch belong to dopamine neurons or MBONs. Our classification data showed that only one output neuron and two input neurons had more than one branch, and most KCs had only one connection with APL neurons (Fig. 3C).

**Fig. 3.**
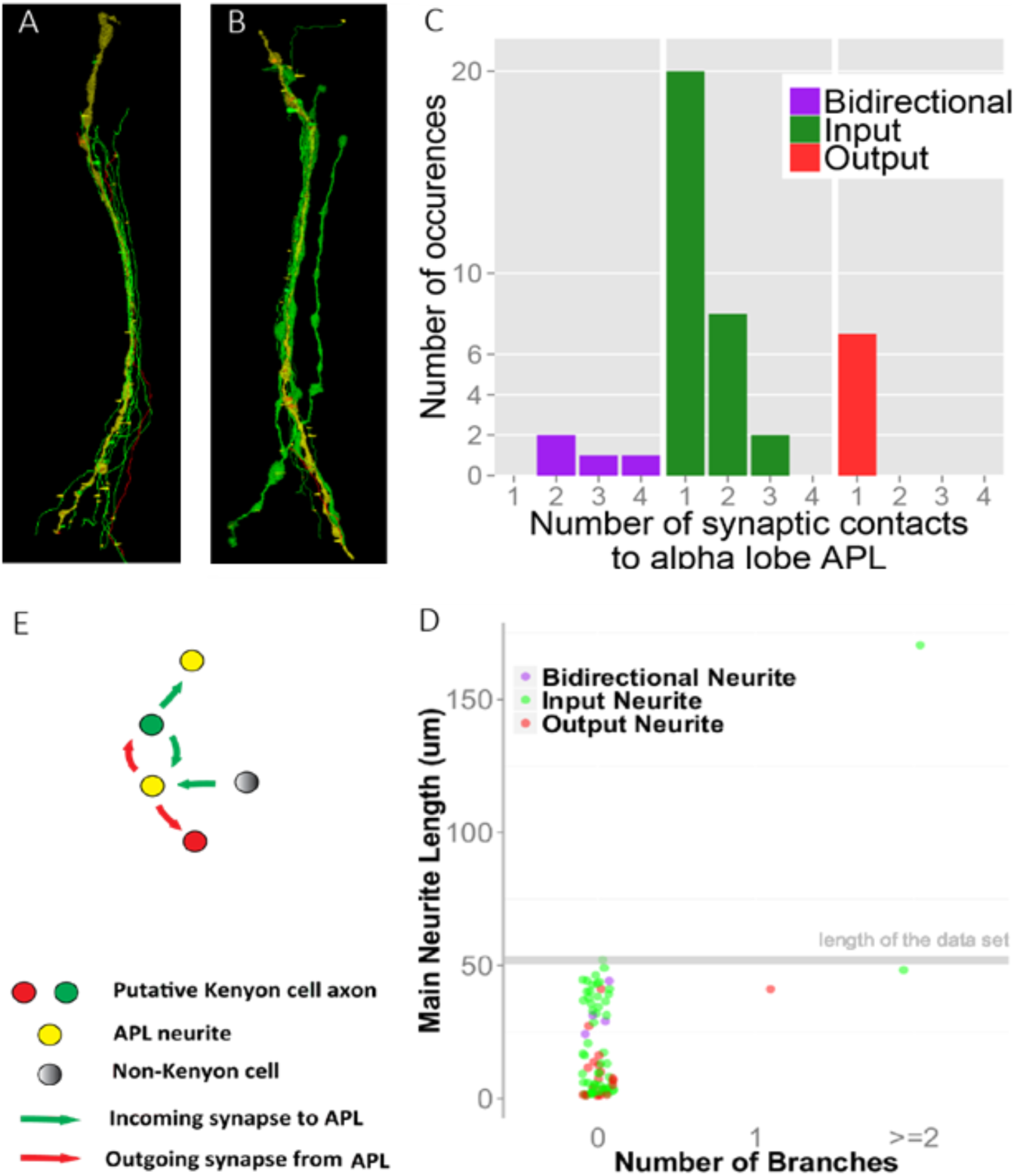
Microcircuit (recurrent network and lateral inhibition) between APL neurons and KCs. **(A)** and **(B)** Two examples of APL input and output neurons. We recognized and traced all of the input and output neurites of APL neurons in the α lobe and the reconstructed 3D structures of some neurites. In this figure, APL neurites are shown in yellow, input APL neurons are shown in green, and APL output neurons are shown in red. The two examples show that these input and output neurons are mostly close to APL neurons; they extend along vertically in a bunch and form synapses with APL neurons, but they separate from the synapses with APL neurons. **(B)** A few reconstructed APL input neurons also have enlarged and thin regions. The APL input synapse is represented in green, the APL output synapse is shown in red, and synapses connecting to other neurons are shown in yellow. **(C)** The numbers of synapses connecting APL neurons with input and output neurons. The green bars indicate the distribution of synapses between APL and input neurons. The red bars indicate the distribution of synapses between APL and output neurons. The purple bars indicate the distribution of synapses between neurons that have both input and output connections with APL neurons. More than 60% of neurons form single synapses with APL neurons. All input and output neurons had a total length greater than 10 µm. **(D)** All neurons connecting to APL neurons are Kenyon-like neurons. The x- and y-axes represent the number and length of the branches, respectively. The morphologies of most input and output neurites contain only one main neurite, i.e., had zero branches. Many neurons with only one main neurite were longer than 25 µm, or even near 50 µm, almost equaling the length of the α lobe in our dataset (52 µm, gray horizontal line in D). Therefore, these neurons nearly extend across the α lobe and have morphologies similar to those of KCs. One neuron with branches extends across the α lobe with multiple lengths of the α lobe (green point in the upper right corner of D). This observation suggests that multiple types of input neurons connect to APL neurons. Green point: Input neurons of APL neurites. Red point: Output neurons of APL neurites. Purple point: Neurons with input and output connections to APL neurites. **(E)** Microcircuits between APL and KC neurons, including recurrent network and lateral inhibition.

We concluded that several circuits exist in the vertical lobes of mushroom bodies between APL neurons and other neurons (Fig. 3E). We found lateral inhibition and a recurrent network between APL neurons and KC neurons in the vertical lobe. This circuit comprising dopamine neurons projecting to KCs constitutes the neural basis for decoding of odor learning, decision making, mutual suppression of relational odor, and reversal learning. EM reconstruction data revealed that in the vertical lobe, KC neurons projected to APL neurons non repetitively (Fig. S11). One strategy for fulfilling the function of specific inhibition is utilizing the local microcircuit between APL neurites and KCs at different compartmental parts with dopamine neurons. Therefore, KC axons may be compartmentally modulated by not only dopamine but also APL neurons.

### APL neurons generate no action potential

Amacrine A17 neurons in which multiple independent microcircuits exist in a single neuron (Grimes et al., 2010), are nonspike and axon-less neurons. Small axon-less neocortical interneurons, which are also nonspike neurons, communicate with each other through dendrodendritic local circuits (le Magueresse et al., 2011). Neurons generating no action potential send signals via graded membrane potential to form microcircuits, whereas neurons generating action potentials function as a single unit; therefore, local circuit is easily formed by non spike neurons but not spike neurons. Other dendrodendritic microcircuits exist between excitatory mitral cells and inhibitory granule cells (Osinski & Kay, 2016). GGNs (the homolog of APL neurons) in locusts normalize the KC odor response via global inhibition, generating no action potentials to odor stimuli (Papadopoulou et al., 2011). Therefore, we examined this electrophysiological property of APL neurons. We depolarized the cell bodies of APL neurons (n=5) with a slope from −30 mV to 80 mV, and none of the APL neurons elicited an action potential (Fig. S12). Such, APL neurons are similar to the axonless amcrine A17 neurons and neocortical interneurons. Moreover, we examined whether electrical stimuli are transferred from APL cell bodies to lobes. Indeed, we failed to record changes in the voltage signal in the lobes upon depolarization of cell bodies.

### APL neurites in the α and α**′** lobe respond to odors independently

First, we tested whether APL neurons in the α and α′ lobes function independently. The reconstructed EM data showed only one neurite connecting the α and α′ lobes in the vertical lobe (Fig. 1FG). The structural separation between the α and α′ lobes of APL neurites indicates that these lobes might function independently based on their local circuitry with KCs in α and α′ lobes separately. Moreover, octopaminergic APL neurites in the α lobe have been shown to maintain anesthesia-sensitive memory, while APL neurites in the α′ lobe facilitate visual reversal learning; this functional separation of the APL – KC at α and α′ lobes circuit is consistent with our hypothesis.

Previous studies found that APL neurons depolarize in response to both odors and electrical stimuli (Liu & Davis, 2009; Wu et al., 2011). APL neurons in the vertical lobe can also respond to odors (Lin et al., 2014). Thus, the different responses property of APL neurons in the α and α′ lobes to odor and electrical stimuli were explored.

To test our hypothesis that APL neurons have independent local circuits in the α and α′ lobes, we used dual-color calcium imaging to observe the responses of APL neurons and mushroom bodies, *GH146-Gal4/UAS-GCaMP6s; 247-LexA/LexAop-R-GECO* (Fig. 4E). Overall, angular distance data showed that the APL neurites responded to odors more generally than did the KCs. These data are consistent with previous research results (Wu et al., 2011). And, APL neurites in the α lobe and α′ lobes were shown to respond to odors differently. Firstly, the maximal APL peak response in the α lobe was much larger than that in the α′ lobe (Fig. 4AB). Additionally, among all of the responsive regions of interest (ROIs), APL neurite at the α lobe tended to respond to more odors than that at the α′ lobe (Fig. 4CD), while they have better odor separation of APL neurite in α′ lobe than in α lobe (Fig. 4EFG). Second, both ON and OFF responses were observed in APL neurites, especially those in the α lobe, which exhibited a greater OFF response, suggesting that KCs have not only ON responses (Li, Li, Lei, Wang, & Guo, 2013) but also OFF responses. Thus, APL neurites in the α and α′ lobes respond to odors differently and independently. This difference might be due to different input KC response properties in the α and α′ lobes.

**Fig. 4.**
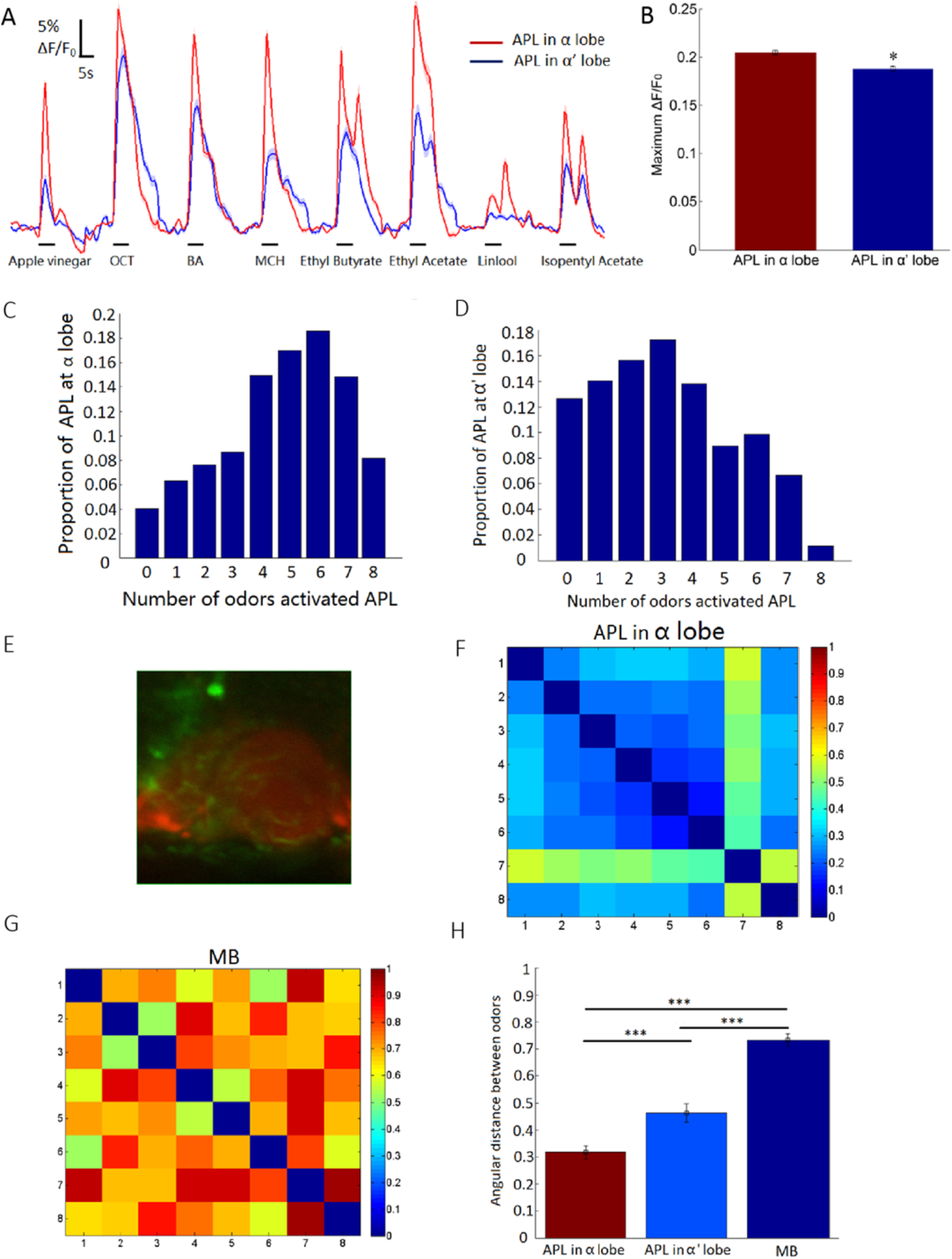
APL neurons in the α and α′ lobes respond to odors differently. **(A)** Mean curves of APL responses to eight odors showing that the peak response of the α lobe is larger than that of the α′ lobe. Odors delivered to the flies included apple vinegar, OCT, BA, MCH, ethyl butyrate, ethyl acetate, linalool, and isopentyl acetate. **(B)** The peak responses of APL neurites in the α lobe are larger than those in the α′ lobe (Wilcoxon rank-sum test, P < 0.05, P = 8.947×10^−6^). **(C)** APL neurites in the α lobe responded to more odor types. **(D)** APL neurites in the α**′** lobe responded to fewer odor types. In (A) and (B), the horizontal axis represents the number of responsive odors, and the vertical axis represents the percentage of responsive ROIs. **(E)** Horizontal map of an APL and mushroom body in the vertical lobe by dual-color calcium imaging. The fly genotype is *GH146-Gal4/UAS-GCaMP6s; 247-LexA/LexAop-R-GECO*, n = 10. **(F)** Angular distance heatmap of APL neurites in the α lobe responding to eight types of odors demonstrating that APL neurites separate odors poorly. **(G)** Angular distance heatmap of a mushroom body responding to eight odors demonstrating that mushroom bodies can separate odors well. **(H)** Mean angular distances of mushroom body and APL responses to different odors showing that mushroom bodies can separate odors better than APL neurites, while APL neurites in the α′ lobe separate odors better than those in the α lobe (Wilcoxon rank-sum test, P < 0.001. P (α & α′) = 0.0013, P (α & MB) = 4.59×10^−17^, P (α′ & MB) = 4.37×10^−8^).

### Odor evokes a local calcium microdomain on a single APL neurite

The calcium signal has been shown to be a good indicator of neuronal activity, especially after the advent of GCaMP6 (Chen et al., 2013). We used a single APL fiber calcium imaging method to observe its calcium signal for different odors (Single neurite imaging, Fig. S13). The alternate arranged enlarged and thin sections of the APL neurites enables the different electrical conductance across the neurite (Rall, 1969). This suggests that a separate calcium activity zone can be seen on the nerve fibers. Secondly, blocking KC output completely abolished the responses of APL neurons to odors physiologically (Lin et al., 2014). Besides, KC codes odor sparsely and separate odor well, only a few KCs respond to more than one odors (Lei et al., 2013). Structurally EM reconstruction data revealed that in the vertical lobe, KC neurons projected to APL neurons nonrepetitively (Fig. S11); Thus, local circuits between APL neurons and KCs enabled the differential and local calcium response of APL neurite to odors.

Therefore, to address whether odors can induce local and different calcium responses in APL neurons, we observed single APL neurite calcium activity at the vertical lobe via expressing GCaMP6 driven by *APL-GAL4*. APL neurites are imaged across layers from top to bottom and then we compared single neurite activities within the same layers, found local calcium microdomains on APL neurites induced by different odor inputs (Fig. 5A). Local calcium domains have been demonstrated to be electrically separated from each other on a neurite (Goldberg, Gabor, Dmitriy, & Rafael, 2003). The imaged APL neurites were classified into three categories: neurites that responded to all odors locally (46%), those that locally responded to some of the eight odors (34%), and those that did not respond to any odors (20%) (Fig. 5ABC) (Table 2). That might due to different APL neurons receive different KCs input at different location. Some KCs could respond to more than one odor, while other KCs code non odor information but temperature and others. These phenomenon were consistent with our hypothesis. Another explanation for the local calcium domains could be an uneven calcium channel distribution. To exclude this possibility, we delivered electrical and odor stimuli separately to Drosophila. The responses to electrical and odor stimuli are first mediated by different neurons; the response to electrical stimuli is primarily mediated by dopamine neurons (Claridge-Chang et al., 2009; Mao & Davis, 2009), which then transfer the information to APL neurons directly or indirectly via KC neurons. In contrast, the response to odor stimuli is mediated by KCs. Second, KC neurons project to APL at different locations. If localized APL neuron responses are due to different inputs and not to an uneven calcium channel distribution, then APL neurons should respond to odors differently when exposed to different stimuli. The inability of these neurons to respond to odor signals activated by an electrical stimulus demonstrates that they are activated by different inputs in different regions. Thus, APL neurites displayed local input-dependent output activity but not a global output signal (Fig. 5D). 3D reconstruction of volume calcium imaging APL neurites via tracing mCherry expression in the vertical lobe showed local calcium responses from the top to the bottom of the lobe (Fig. 5E). Additionally, responses of the same neurites across several layers demonstrated significant local responses to odors (Fig. 5F). These local calcium domains suggest poor propagation of the calcium signal along the APL neurite. This local calcium response property is common on divided leaves of the vertical lobe (Fig. 5G). We also calculated the preference of odors in different APL neurites, which displayed a different preference but not an even response (Fig. S14).

**Fig. 5.**
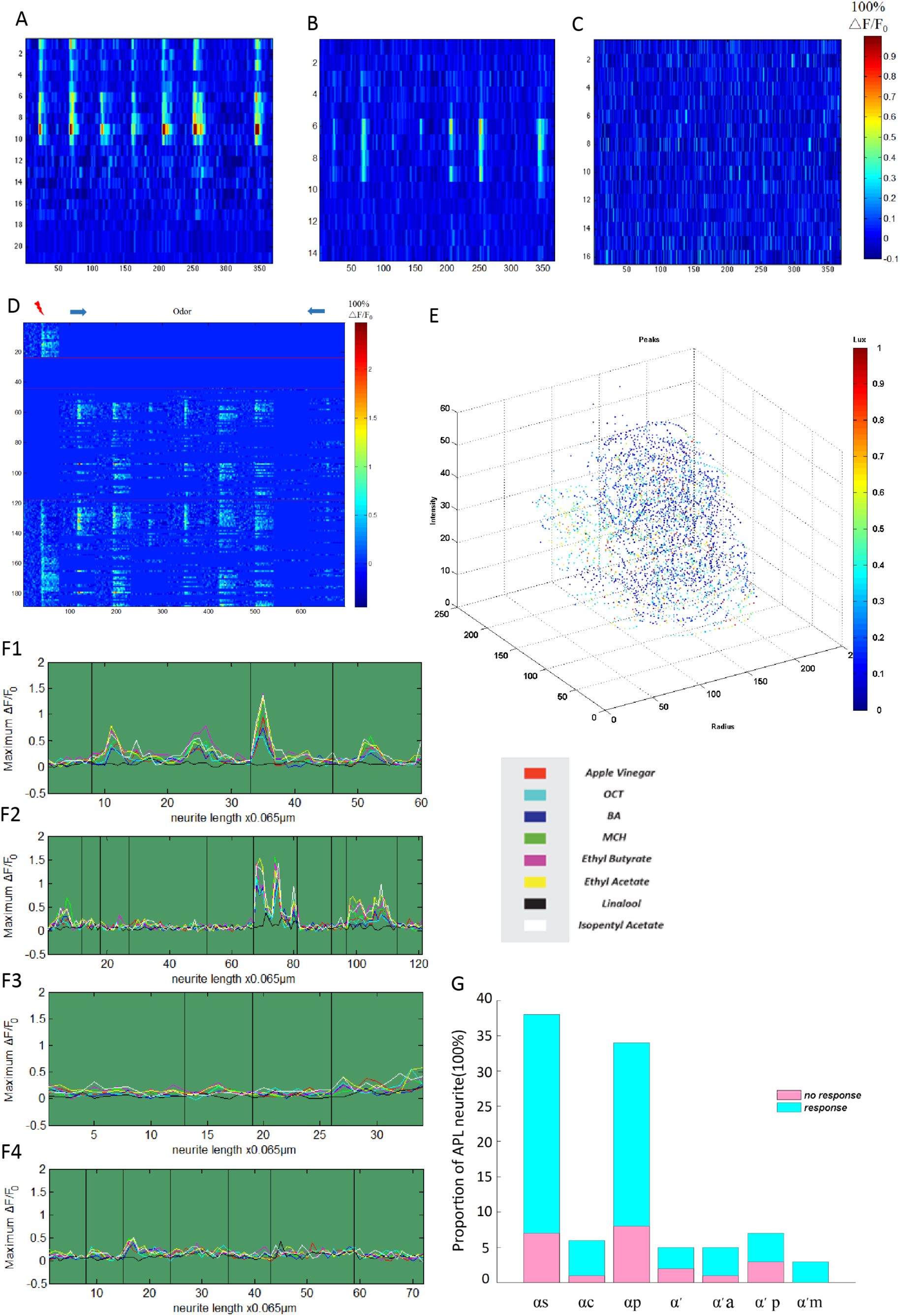
Localized calcium domain on a single APL neurite. **(A)** APL neurites that respond to some odors (odor stimulus label is the same as in Fig. S13). **(B)** APL neurites that respond to all odors. **(C)** APL neurites that respond to no odors. In (A), (B) and (C), the vertical axis represents the length of neurites, the horizontal axis represents the time of calcium imaging with odor stimuli, and the heat value represents the peak responses to odors. Eight different odors were delivered in the sequence listed in Fig. 4A for each heatmap. *GH146-Gal4/UAS-GCaMP6s; UAS-mCherry/+*, n = 5. **(D)** Electrical stimuli activated different APL neurites than odor stimuli. The first area represents APL neurites activated by only electrical stimuli. The second area represents APL neurites that did not respond to either odor stimuli or electrical stimuli. The third area represents APL neurites that only responded to odor stimuli. The fourth area represents APL neurites responded to both odor stimuli and electrical stimuli. The red lightning bolt represents a 60 V electrical stimulus applied for 5 s. Eight different odors are shown in Fig. 4A. *GH146-Gal4/UAS-GCaMP6s; UAS-mCherry/+*, n = 3, n (ROI) = 190. **(E)** Local peak responses to apple vinegar in the 3D structural representation of APL neurites. The x- and y-axes represent the horizontal level, the z-axis represents the vertical level, and the light intensity represents the response intensity at that point. **(F)** The same APL neurite across different layers demonstrating a localized response property to odors. F1-F4 represent four different APL neurites. The horizontal axis represents the length of APL neurites across different layers. Each black line represents the joint between two neurites at different layers belonging to the same neurite. The vertical axis represents the peak response of calcium imaging to odors. *GH146-Gal4/UAS-GCaMP6s; UAS-mCherry/+*, n = 4. **(G)** APL neurites in different subareas of the mushroom body vertical lobe, all displaying the local response property. The horizontal axis represents APL neurites in different subareas of the mushroom body, and the vertical axis represents the percentage of responding (blue bar) and nonresponding APL neurites (pink bar).

**Table 2.** Statistical data of various APL neurite responses to odors

| **Kinds of APL neurite responses to 8 odors** | **Proportion** |
| --- | --- |
| Local response to 8 odors | 47/103 (46%) |
| Non response to odors | 21/103 (20%) |
| Local response to part odors | 35/103 (34%) |

### APL neurites send out signals locally both in the calyx and the vertical lobe depending on odor stimuli

Because APL neurites respond to odors locally depending on the local input in the vertical lobe, we examined whether they also send out signals locally in the calyx and vertical lobe, since the same local circuits are present in both the calyx (*unpublished*) and vertical lobe. We expressed a presynaptic-specific calcium indicator, syt-R-GECO, driven by *GH146-Gal4* and *APL-Gal4* and then observed APL output signal responses to 8 different odors in the vertical lobes (Fig. 6A) and calyx. Heatmaps of APL neuron presynaptic calcium activity signals displayed local odor dependent activity in both the vertical lobe and calyx (Fig. 6BC).

**Fig. 6.**
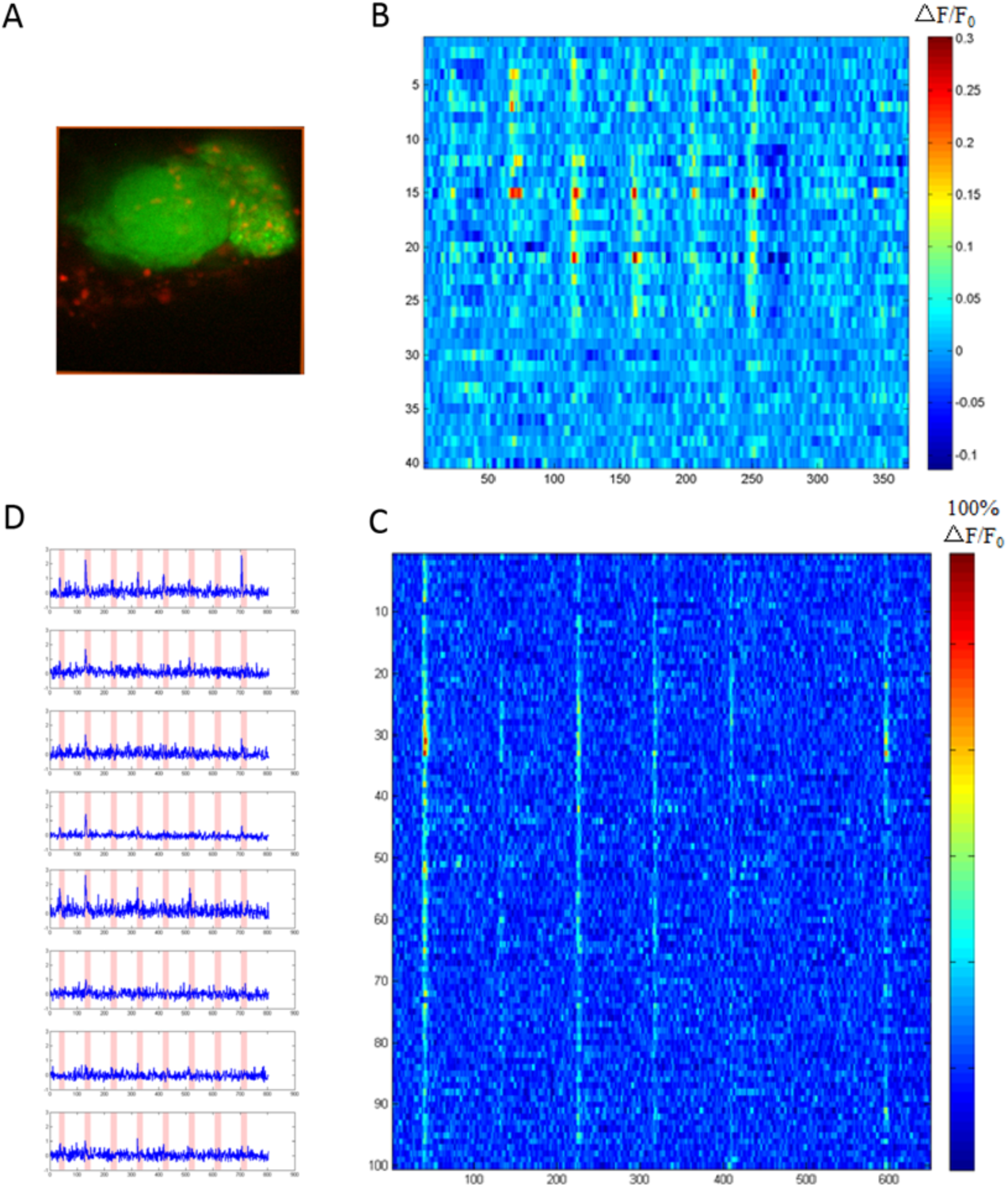
Localized responses of an APL presynaptic calcium indicator to odors. **(A)** Distribution of presynaptic APL neuron proteins in the mushroom body vertical lobe. *GH146-Gal4/+; UAS-syt-R-GECO/247-LexA-LexAop-GCaMP6s*, n = 7, n (ROI = 40). The green area represents labeled KCs, and the red area represents presynaptic proteins in APL neurons. **(B)** Local responses of an APL presynaptic calcium indicator to eight odors in the mushroom body vertical lobe. **(C)** Local responses of an APL presynaptic calcium indicator to eight odors in the mushroom body calyx. **(D)** Eight ROIs of APL-localized presynaptic site responses to 8 different odors in the calyx. (C and D) *UAS-syt-R-GECO/+; APL-Gal4/+*, n = 3, n (ROI) = 100. (BCD). Odor stimuli are the same as mentioned in Fig. 4A.

In the calyx, regions of APL neurites labeled by presynaptic proteins responded broadly to the first odor, providing structural evidence of global APL neurite distribution (Fig. 6C). In contrast, only some presynaptic protein-labeled APL neurite regions responded to other odors, and the activated presynaptic APL neurite areas differed among odors, demonstrating that APL neurites show local but not global odor-dependent activity. Examples of a single presynaptic calcium unit displaying odor-specific local presynaptic activity are shown in Fig. 6D.

Previous studies found that GABAergic neurons project to KC dendrites and projection neuron axons in the calyx (Yasuyama, Meinertzhagen, & Schurmann, 2002), implying that a feed-forward inhibitory circuit between APL neurons and projection neurons may exist in the calyx (Olsen & Wilson, 2008). Whether KCs or projection neurons project to APL neurons in the calyx, forming local neural circuits with APL neurons, requires further study.

In summary, the evidence presented support the presence of a local recurrent circuit of APL neurons and KCs in the vertical lobe (Fig. 7).

**Fig. 7.**
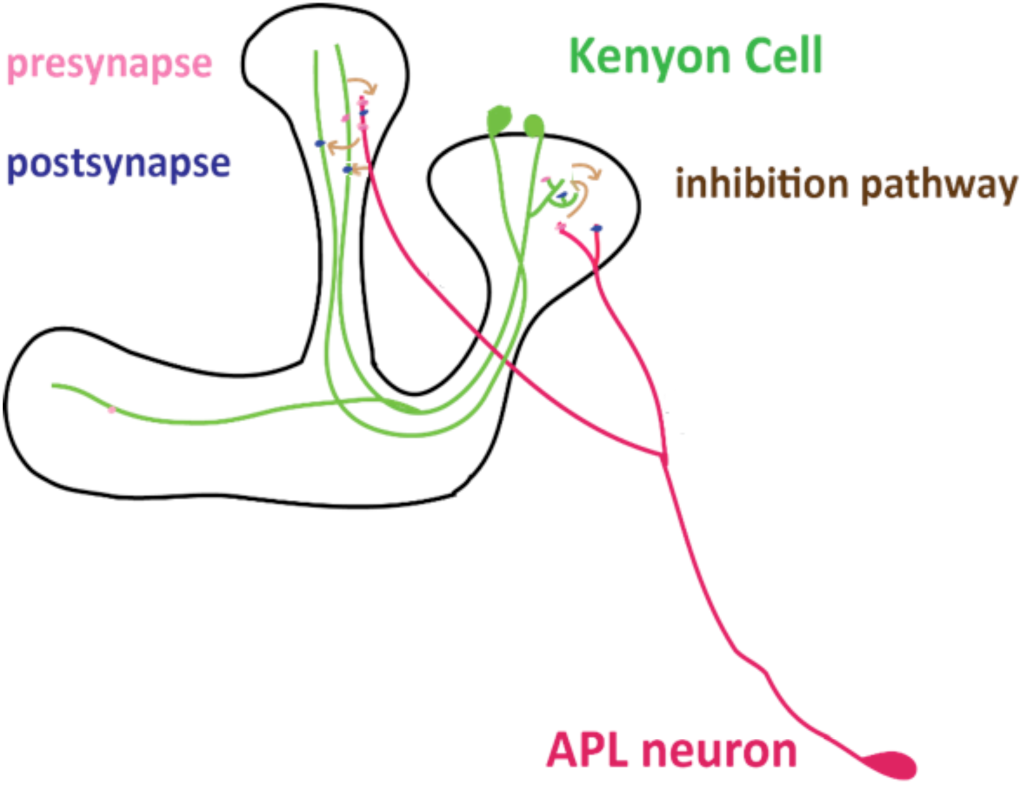
Working model of the circuit between APL neurons and KCs. Mutual and local inhibition circuits between APL neurons and KCs in the mushroom body vertical lobe. We hypothesize that local circuits exist in the mushroom body calyx. The green curves represent KCs, and the red curves represent APL cells. Connection circuits that potentially exist between other neurons and APL cells are not shown.

## Discussion

Mushroom body and APL neurons play important roles not only in learning and memory but also in high cognitive behavior, reversal learning, learning, and decision making (*unpublished*). The circuit of APL and KCs in the mushroom body underlies this behavior in reversal learning. To reveal the specific circuits that have remained unknown, we applied series continuous EM methods. Here, we reconstructed the connections between the mushroom body and APL neurons in the vertical lobe of the mushroom body via FIB-SEM and genetic labeling of APL neurons with HRP. We found recurrent network and lateral inhibition connections between APL neurons and KCs in the vertical lobe of the mushroom body. Reconstruction of the mushroom body in *Drosophila* larva also found APL-KC recurrent networks (Eichler et al., 2017), which are the neural basis underlying high cognitive behavior. Other reconstruction neural circuit work in the calyx of the mushroom body also revealed a recurrent network between the APL neurons and KCs (Zheng et al., 2018). Moreover, while dopamine neurons projecting to mushroom body axons result in local KCs function, APL neurons receive input from KCs and also well intermixed with mushroom body compartmental structure, whether APL neurons function locally remained uncertain. APL neurons exhibited intermixed presynapses and postsynapses in both the calyx and vertical lobe. We thus used EM reconstruction, electrophysiology and calcium imaging methods to explore the local function of APL neurons. In the vertical lobe, APL neurites resembled a string of beads with input synapses and output synapses intermixed on enlarged and thin sections of the neurite. KCs were found to project to APL neurons nonrepetitively (Fig. S11), forming a local circuit. Furthermore, using a single neurite calcium imaging method, we found that local calcium domains enabled electrical separation of the APL neurite. APL neurons were found to send odor-dependent signals in a local but not global manner not only at calyx but also at lobe. Furthermore, the vertical lobe was shown to have odor-dependent properties but not global properties. At the vertical lobe, APL neurites only formed connections with 2/3 KC neurons, which is insufficient for a global inhibition network. Consistent with the functional separation of the α lobe and α′ lobe in anesthesia-sensitive memory, Reconstructed APL neurons exhibited a separation at these lobes and APL neurons exhibited independent responses to odors in these lobes. These findings support the hypothesis that APL neurons function locally.

To reconstruct brain circuits on the nano scale, two strategies in volume reconstruction have been applied till now: one is dense reconstruction of all neurons in a specific area and the other is sparse reconstruction, reconstruction of sparsely labeled neurons and their connected neurons. The mushroom body of *Drosophila* larva (Eichler et al., 2017), the calyx of adult *Drosophila* (Zheng et al., 2018), and the entire brain structure of *C elegans* (Cook et al., 2019) have been reconstructed using the first method. The second method has been pioneerly applied in our work. Due to its specific labeling, the second method takes priority in dramatically less time and labor consumption of data acquisition, analysis and data storage than the first one. Since the first one needs to get the whole brain EM data, and then focus on specific region to reconstruct all complete neuron structure, the second one only get the specific brain EM data and reconstruct the labeled neurons and its connection neurons. Due to these conveniences and accuracy of sparse reconstruction, reconstruction work of specific interesting brain area neuron circuits with few group of people could realize and be valuable. To approach this aim, we have first overcomed the technique challenge of specific labelled EM sample preparation, continuous labeled neurons and better image contrast. The HRP labeled method, which limitation is relative small neural network boundary, is more reliable and highly permeable in all published methods (Larsen, 2003; Shu et al., 2011) after we tested. Besides, in more specific brain area EM acquisition of Drosophila and C elegans, higher precision imaging method dual beam series scan electron microscopy (FIBSEM) (Heymann et al., 2006; Knott, Marchman, Wall, & Lich, 2008), which gives much more reliable EM data because its thinnest imaging layer than serial section TEM (ssTEM) (Zheng et al., 2018), serial scanning TEM (ssTEM) (Briggman & Bock, 2012), automated tap-collecting ultramicrotome SEM (ATUIM-SEM) ((Hayworth, Kasthuri, Schalek, & Lichtman, 2006) and serial block-face SEM (SBSEM) (Denk & Horstmann, 2004) is applied.

APL neurons and mushroom bodies play important roles in highly cognitive behavior, including visual reversal learning, olfactory reversal learning and visual decision making. Direct evidence suggests that the connection between APL neurons and KCs participates in the reversal learning behavior of *Drosophila* (Wu et al., 2012). However, the specific neural circuit between APL and KC neurons has remained undiscovered. Our EM work in the vertical lobe of the mushroom body identified two main circuits: a recurrent network and a lateral inhibition circuit. They might underlie these behaviors. Furthermore, simulation work on recurrent networks has demonstrated that these networks are able to facilitate a winner-take-all neuronal signaling process in limited amounts of time (Liu, Dang, & Cao, 2010) and will quickly realize adaptive orthogonal filters and associative memory (Kohonen & Oja, 1976). Indeed, recurrent networks in the cortex are important in action selection (Haga & Fukai, 2018), decision making (Wang, 2008; Wong, 2006), learning (Gianluigi, Amit, & Nicolas, 2003) and reversal learning processes. The mushroom body of *Drosophila* and the cortex of mammals have been suggested to be evolutionarily identical (Tomer, Denes, Tessmar-Raible, & Arendt, 2010). Thus, the recurrent network identified here may be evolutionarily conserved.

APL neurons in the vertical lobe of the mushroom body display local activity to odor stimuli. Previous work showed that APL neurites might not have strict local circuits; upon selective blockade of different KC subtypes, there were no significant increases in the responses of the corresponding KCs (Lin et al., 2014). This phenomenon could also be explained by the existence of a connection between different KC subtypes, potentially compensating for the effect of blocking a single KC subtype. Indeed, recent studies have identified a gap junction between different KC subtypes, but whether a chemical connection exists between different KC subtypes remains to be explored. Therefore, local APL circuitry could also exist under this condition. Another possibility is that DPM neurons in different lobes might be well connected such that even if one KC subtype is blocked, DPM neurons connected with other subtypes might compensate for this effect due to their ability to elicit the release of acetylcholine from KCs via gap junctions between APL and DPM neurons. However, more experimental evidence is needed to support this possibility. The third possibility is that extrinsic inhibitory interneurons may broadly receive input from KC axons and deliver output to APL neurons. Another work showed that APL neuron activity was global or specific depending on the olfactory input strength to the mushroom body (Inada, Tsuchimoto, & Kazama, 2017). In the calyx, partial activation of projection neurons induces partial KC inhibition, which arises from the local but not global activity of APL neurons. This finding is consistent with our finding in the calyx. The calcium response of APL neurons throughout the horizontal lobe was found to increase in response to increasing odor concentration gradients. Increased odor concentration will increase KCs response, which induce high APL response property, it does not mean that the response is global. First, the increase in the APL neuron response cannot exclude the possibility of a DPM neuron electrical coupling effect (Pitman et al., 2011; Wu et al., 2011). More importantly, the difference in research methods used, such as whole-area imaging versus single neurite imaging, might have led to different conclusions. The single neurite calcium imaging method used here provides a more precise signal than the previous method. Another possible explanation is that APL neurites in the horizontal lobe might function differently from those in the vertical lobe.

Local circuits between APL neurons and KCs in the vertical lobe function in compartmental functional mushroom body structures. Mushroom bodies contain 15 independent compartmental function structures separated by locally projecting dopamine and extrinsic neurons. This functional compartmentalization increases the computation capacity of mushroom bodies (Aso et al., 2014a), and familiar strategies are applied by amacrine A17 neurons (Grimes et al., 2010). Broadly innervated APL neurons also work locally, which is another strategy to increase the computational complexity in a limited space. The local circuits in APL neurons act together with the compartmental structures of mushroom bodies and facilitate different mushroom body area responses, and similar local circuits have been identified in amacrine A17 neurons. Additionally, our data suggest that APL neurons exhibit segmental plasticity, which is also supported by evidence that APL neurons specifically respond to learned odors (Liu & Davis, 2009). The distance between local circuits in APL neurons and whether these circuits are restricted to thin and enlarged regions, or even beyond these regions, remains unanswered. This question needs to be further studied by depolarizing a single APL neuron input and then observing the distance of signal propagation (Grimes et al., 2010). This technology is difficult to realize due to the fleeting calcium signal on a single neurite. More questions including what’s the relationship of APL local neurons and DA-MBON-KC compartmental structure also remains to be further study. And what’s the contribution of APL and DA separately on local KC signal.

Another circuit found between APL neurons and KCs is a lateral inhibition circuit. Lateral inhibition between APL and KCs increases the signal contrast. The circuits might not only mediate competition between different types of odor information but also other sensory information in mushroom bodies, as mushroom body KCs encode not only olfactory information but also visual information, taste (Kirkhart & Scott, 2015) and temperature (Frank, Jouandet, Kearney, Macpherson, & Gallio, 2015). However, to complete understand the whole circuit function, the identity and role of other neurons connected with APL neurons remains to be explored in future work. As mentioned previously, dopamine neurons form dopaminergic connections with APL neurons. Together, dopamine neurons (Ke, Guo, Peng, Wang, & Aike, 2007), KCs and APL neurons form the salience-based decision-making circuit (Guo, Zhang, Ren, Su, & Chen, 2016). The specific circuit remains unexplored, and its identification may enhance our understanding of decision-making processes. Dopamine may also act with APL neurons and KCs to effect reversal learning behavior. Here, dopamine encodes value information about KCs, and MBONs encode the decision-making behavioral output? Additionally, the 15 functionally distinct compartments of mushroom bodies are suggested to form a multilayered feedforward learning module in which dopamine encodes reward and APL neurons encode punishment. However, the specific circuit for this coding and computing behavior requires more detailed information (Aso et al., 2014a). Finally, whether decision-making circuits also include different compartmental parts of the mushroom body remains to be explored.

## Methods

### Fly strains

The flies used in this experiment were as follows: *APL-GAL4* (Ann-Shyn Chiang), *GH146-Gal4* (Tanaka et al., 2008), *247-LexA*, *UAS-HRP-mCD8-GFP* (Jean-Paul Vincent), *UAS-GCaMP6s*, *UAS-syt-R-GECO (unpublished)*, *LexAop-GCaMP6s*, *UAS-synaptobrevin-GFP* (Bhattacharya et al., 2002), *UAS-mGFP*, *UAS-n-synaptobrevin-GFP* and *UAS-mCherry*. All flies were raised in an incubator at a constant temperature and humidity of 25°C and 50%, respectively, with a 12 h dark/12 h light circle. The flies were supplied Bloomington standard food, and crossed flies were collected as virgins for calcium imaging experiments.

### EM tissue preparation

Three-day-old male *GH146-Gal4>UAS-HRP-mCD8-GFP* flies were used for EM preparation. The *Drosophila* brains were dissected according to custom procedures. Fixation was performed to ensure that the *Drosophila* brains were HRP-positive under a light microscope. The brains were fixed in glutaraldehyde for 30 min at 25°C, dyed with HRP, and then washed with 0.1 M PB three times for 5 min each time and incubated with TSA biotin for 16 h at 4°C. Then, the brains were washed and incubated with ABC for 16 h at 4°C. The brains were then washed with Tris-HCl, incubated with DAB first at 4°C for 30 min and then at 25°C for 15 min, and washed with 0.1 M PB. For EM sample preparation, the brains were fixed with prechilled 2% OsO4 PB for 2 h at 4°C, washed with ddH2O, dyed with 2% uranyl acetate for 20 h at 4°C, washed with ddH2O, and then pre-embedded in 1% agarose at 42°C. After the agarose set, it was trimmed into a trapezoid. Dehydration and resin impregnation were performed according to the standard EM method. Fly brain resins embedded in agarose were dried in an incubator at 37°C for 12 h, 42°C for 12 h and 60°C for 48 h. The resins were carefully trimmed to expose the vertical lobe of the mushroom body for dual beam electron microscope imaging.

### EM data imaging and reconstruction procedures

#### Imaging data capture parameters and results

We performed carbon plating for the EM samples and glued them to the EM sample plate. We then placed an EM sample with plates inside the dual beam scanning electron microscope (Helios Nano Laboratory 600i, FEI). A 1.5-µm-thick layer of platinum was placed onto the surface perpendicular to the block face to be imaged. A focused electron beam was used to image the block face after the focused ion beam was used to remove a 40-nm-thick block face and expose a new one to be imaged. These two steps were repeated until the entire dataset was finished. Platinum plating and imaging capture were performed using AutoSlice & View^TM^ software (FEI). In total, 1300 images were acquired, and every image pixel was 4096×4096 with a pixel distance of 7 nm.

#### Reconstruction procedure (neurite reconstruction: node, line, 3D reconstruction)

All images were aligned in FIJI TrackEM2 by SIFT alignment. Due to the lower accuracy of autosegmentation of neurites, manual labor was preferred. Approximately 5-6 annotators separately identified darkened neurites and synapses in each layer and then cooperated to proofread and vote on the merged results according to a strict standard. Only nodes or synapses more than 80% positive were subjected to a recheck, and only 100% positive data were accepted to ensure the authenticity of the dataset. Additionally, the missed data (non-100% positive data) were added in the next analysis procedure. Then, the results from all of the annotators were merged for further analyses. The overall marking/proofreading-voting-merging cycle was repeated with the help of modified TrackEM2. In all analyses, all neurons and synapses were checked at least 3 times, and several areas were checked 5 times. In the end, we are confident that we obtained 100% correct data. In addition, based on this process, we annotated APL neurite skeletons, APL synapses, input neuron skeletons, input neuron synapses, output neuron skeletons and output neuron synapses one-by-one. To obtain volume data for some neurites, we performed neurite segmentation by a semiautomatic method (*Supplemental material)*.

#### Neurite radius extraction and data analysis

Neurite radius computation was based on 3D neurite data. The minimum section area (A) of a neurite at a specific node was calculated, and the radius of a specific node was calculated as the radius of a minimum section area. Thus, 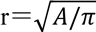. The code for neurite radius computation was written using Python (v2.7, www.python.org), and the results were saved as xml files. All other data were acquired by running scripts in FIJI via the TrackEM2 program interface. We then analyze these data in R and plotted the data via ggplot2. The data were modified, and the typeset figures were created using an illustrator (Adobe Systems Inc.).

### In vivo calcium imaging

#### Odor delivery

Eight different odorants, 4-methylcyclohexanol (MCH), octanol (oct), butyl acrylate (BA), apple vinegar, ethyl butyrate, ethyl acetate, linalool, and isopentyl acetate, were separately diluted in mineral oil at a ratio of 1:100. All odorants were purchased from Sigma at the highest purity possible. The odorants were pumped into the fly chamber at a final speed of 400 ml/min via a pipe (microdiaphragm pump, model HC400DC). The ventilated pipe was placed 1 cm from the fly antenna. The time (5 s) and interval (18 s) of odor delivery were controlled using MATLAB software. The odor delivery timing accuracy was adjusted using a photoionization detector (miniPID, Aurora Scientific Inc.). All of the above odorants were purchased from Sigma-Aldrich at the highest purity possible.

#### Electrical stimuli

Flies were prepared as mentioned before for in vivo calcium imaging. Then, the fly was placed in a chamber with copper on it, which was connected to electrical power, ensuring that their abdomens and legs touched the copper. The time and duration of electrical stimuli were controlled by MATLAB software. Finally, 60 V of electrical power was delivered to the fly for 5 s.

#### Calcium imaging

Flies expressing *UAS-GCaMP6s* via *GH146-Gal4* were anesthetized on ice and then transferred to 4°C. They were then placed on a classic imaging plate with their antenna beneath the plate. The point at which the fly and plate connected was covered with ultraviolet glue (UV 353 Loctite) and then fixed with an ultraviolet lamp for 20 s. The head cuticle between the two eyes, fat body and trachea were carefully removed with fine forceps. The fly brain was immersed in extracellular solution for cells (ESC) (Wilson, Turner, & Laurent, 2004). The plate was then reversed, and the proboscis cuticle was cut to ensure that the skeleton was bendable and stable. The reversed plate was then placed under a NIKON FN1 microscope with a Nikon Apo LDW 25x/1.1w water objective. The excitation wavelength was 488 nm. We acquired images with 256×256 pixels at a speed of 8 Hz. For multiple-layer imaging, each step was 4 µm. The setup for the experiment, including odor stimuli and electric stimuli, is shown in Fig. S15.

Dual-color flies containing *GH146-Gal4/UAS-syt-R-GECO*; *247-LexA-LexAopGCaMP6s* and *GH146-Gal4/UAS-GCaMP6s*; *UAS-mCherry/+* and *GH146-Gal4/UAS-GCaMP6s*; and *247-LexA/LexAop-R-GECO* were imaged with an additional excitation laser at 568 nm. Other flies included *UAS-syt-R-GECO/+; APL-Gal4/+*.

#### Imaging data analysis

We separately aligned the imaging data channels using scripts or TurboReg in FIJI. We then drew continuous and even 2-µm ROIs along the neurite. Calcium changes in the ROI were calculated according to the odor delivered. We performed multilayer neurite reconstruction using TrackEM2 in FIJI. The analysis described below was performed in MATLAB. We identified the average calcium signal at 9 s before the odor or electrical stimulus was applied as F_0_ and the calcium change as (F- F_0_)/F_0._ The peak response 3 times larger than the standard deviation before odor delivery was selected as the responsive ROI, and other points were set to zero. Line traces of APL neurons to different odors represent the mean calicum response plus s.e.m., and statistical analyses were conducted with the Wilcoxon rank-sum test.

The angular distance separation between mushroom bodies and APL neurons for eight different odors was calculated as follows: the peak response of all ROIs (KCs or APL neurons) to one odor represents a vector, and the difference between two vectors was measured by the angle θ between two response vectors (Lei et al., 2013). Thus, the angular distance separation of two odor representations was calculated by 1-cosθ, which ranges from 0 to 1. The average of 28 pairs of odor separation between 8 odors was calculated to measure the separation abilities of KCs and APL neurons. The angular distance separation of different regions of APL neurites was used as the peak response to 8 odors as a vector.

### Electrophysiology

Flies were prepared as previously mentioned, with the brain exposed to the air, stably fixed in the chamber, and immersed in ESC. Whole-cell recordings were performed under visual control using an Olympus BX61WI microscope with a 40× water immersion objective. The recordings were performed at 20–25°C with a MultiClamp 700B amplifier (Molecular Devices). The data were low-pass filtered at 10 kHz and acquired at 20 kHz with a Digidata 1440A digitizer (Molecular Devices).

Fly brains were dissected and immersed in an extracellular saline solution in a small dish. The saline composition was as follows: 103 mM NaCl, 3 mM KCl, 5 mM N-Tris (hydroxymethyl) methyl-2-aminoethane-sulfonic acid, 10 mM trehalose, 8 mM glucose, 26 mM NaHCO_3_, 1 mM NaH_2_PO_4_, 1.5 mM CaCl_2_, and 4 mM MgCl_2_ (adjusted to 275 mOsm). The saline was gassed with 95% O_2_/5% CO_2_ to a final pH of 7.3. The sheath covering the target neuron was removed with fine forceps. *GH146-GAL4* was used to label APL neurons. Patch-clamp electrodes (10–20 MΩ) were filled with the following solution: 140 mM potassium aspartate, 10 mM HEPES, 1 mM KCl, 4 mM MgATP, 0.5 mM Na_3_GTP, 1 mM EGTA and 1% biocytin hydrazide (pH 7.3, adjusted to 265 mOsm). In most of the recorded neurons, a small constant hyperpolarizing current (−50–0 pA) was injected to maintain the membrane potential between −60 and −50 mV. All reagents were purchased from Sigma-Aldrich unless stated otherwise. The fly genotype used in this experiment was *GH146Gal4/UAS-tdtomato; 247-LexA/LexAop-NpHr*, n=5.

### Immunostaining

Immunostaining was performed as previously reported (Wu & Luo, 2006) with the following modifications: 4-day-old female adult brains were dissected in PBS and then fixed in 4% (wt/vol) paraformaldehyde (Sigma-Aldrich) in PBS on ice for 20 min. After three washes (20 min each) in 0.3% (vol/vol) Triton X-100 (Sigma-Aldrich)/PBS, the samples were blocked with 5% (vol/vol) normal goat serum (NGS, Vector Laboratories) in 0.3% (vol/vol) Triton X-100/PBS for 1 h. The samples were then incubated with a primary antibody diluted in 5% NGS in 0.3% (vol/vol) Triton X-100/PBS at 4°C overnight. After three washes (20 min each) in 0.3% (vol/vol) Triton X-100/PBS, the samples were incubated with a secondary antibody diluted in 5% NGS in 0.3% Triton X-100 at 4°C overnight. After three washes (20 min each) in 0.3% (vol/vol) Triton X-100/PBS, the samples were mounted with SlowFade Gold antifade reagent (Invitrogen Molecular Probes). The primary antibodies included mouse monoclonal nc82 [1:100; Developmental Studies Hybridoma Bank (DSHB)] and rabbit serum anti-GFP (1:500, Invitrogen Molecular Probes). The secondary antibodies included Alexa Fluor 568 goat anti-mouse IgG (1:200, Life Technologies) and Alexa Fluor 488 goat anti-rabbit IgG (1:200, Life Technologies).

For imaging, Z-stacks of the brains were acquired on an Olympus FV1000 system with a 60x oil immersion objective at 4× magnification. The volume resolution (xyz) of the mushroom body vertical lobe was 0.077×0.077×0.250 µm^3^.

## Acknowledgements

We thank Ann-Shyn Chiang for kindly offering *APL-GAL4* to us, thank Jean-Paul Vincent for offering *UAS-HRP-mCD8-GFP* lines. We thank Aike Guo (Laboratory of Learning and memory, Institute of Neuroscience, CAS) for his great mentor work, kindly and generously financial support of this work, and Ke Zhang for his assistance and support. Thank Ji Gang, JianGuo Zhang for EM support (Laboratory of EM, Institute of Biophysics, CAS), Qian Hu for Confocal light microscopy support (Laboratory of Optical Imaging, Institute of Neuroscience, CAS), Hua Han and LiJun Shen for FIJI modification (Laboratory of Microreconstruction, Institute of Automation, CAS). Li Hui for trial of artificial recognition and automatic recognition of neurons (Fudan University). Li Qiang for computing trials for automatic recognition and tracking (Fudan University). Thank ZhengChang Lei for the productive discussion. This work was supported by 973 Program Grant 2011CBA00400 (to A.G.); Natural Science Foundation of China Grants 30921064, 90820008, and 31130027 (to A.G.); and Strategic Priority Research Program of the Chinese Academy of Sciences Grant XDB02040100 (to A.G.).

## Competing interest

The authors have no conflicts of interest to declare.

## Author contributions

Guangxia Wang designed and performed the calcium imaging experiments, wrote the MATLAB code, analyzed and displayed the calcium imaging and electrophysiological data, and wrote the manuscript. BangYu Zhou and GuangXia Wang designed the EM experiments and synapse-counting experiments and performed the EM experiments. BangYu Zhou and GuangXia Wang worked with ShengXiong Wang, KaiYang, and JianJian Zhao on the reconstructed 3D APL EM connectome data as a team. BangYu Zhou extracted and displayed EM data information. Xing Yang performed the electrophysiology experiment. YiMing Li performed the calcium experiment on the APL output signal in the calyx. LiJun Shen modified the FIJI software. Kai Yang and ShengXiong Wang performed the light microscopy synapse-counting experiment. There is no conflict of interest in this project.

## Notes

#### Summary of Updates

Figure 2,3,6 and Fig.S6,S7,S14 revised

## References

Aso, Y., Hattori, D., Yu, Y., Johnston, R. M., Iyer, N. A., Ngo, T. T., … Rubin, G. M. (2014a). The neuronal architecture of the mushroom body provides a logic for associative learning. eLife, 3, e04577. doi:10.7554/eLife.04577

Aso, Y., Sitaraman, D., Ichinose, T., Kaun, K. R., Vogt, K., Belliart-Guerin, G., … Rubin, G. M. (2014b). Mushroom body output neurons encode valence and guide memory-based action selection in *Drosophila*. eLife, 3, e04580. doi:10.7554/eLife.04580

Bhattacharya, S., Stewart, B. A., Niemeyer, B. A., Burgess, R. W., McCabe, B. D., Lin, P., … Schwarz, T. L. (2002). Members of the synaptobrevin/vesicle-associated membrane protein (VAMP) family in *Drosophila* are functionally interchangeable *in vivo* for neurotransmitter release and cell viability. Proceedings of the National Academy of Sciences of the United States of America, 99(21), 13867–13872. doi:10.1073/pnas.202335999

Briggman, K. L., & Bock, D. D. (2012). Volume electron microscopy for neuronal circuit reconstruction. Current Opinion in Neurobiology, 22(1), 154–161. doi:10.1016/j.conb.2011.10.022

Chen, T.-W., Wardill, T. J., Yi, S., Pulver, S. R., Renninger, S. L., Amy, B., … Vivek, J. (2013). Ultrasensitive fluorescent proteins for imaging neuronal activity. Nature, 499(7458), 295–300. doi:10.1038/nature12354

Claridge-Chang, A., Roorda, R. D., Vrontou, E., Sjulson, L., Li, H., Hirsh, J., & Miesenbock, G. (2009). Writing memories with light-addressable reinforcement circuitry. Cell, 139(2), 405–415. doi:10.1016/j.cell.2009.08.034

Cohn, R., Morantte, I., & Ruta, V. (2015). Coordinated and compartmentalized neuromodulation shapes sensory processing in *Drosophila*. Cell, 163(7), 1742–1755. doi:10.1016/j.cell.2015.11.019

Cook, S. J., Jarrell, T. A., Brittin, C. A., Wang, Y., Bloniarz, A. E., Yakovlev, M. A., … Emmons, S. W. (2019). Whole-animal connectomes of both *Caenorhabditis elegans* sexes. Nature, 571(7763), 63–71. doi:10.1038/s41586-019-1352-7

Denk, W., & Horstmann, H. (2004). Serial block-face scanning electron microscopy to reconstruct three-dimensional tissue nanostructure. PLoS Biology, 2(11), e329. doi:10.1371/journal.pbio.0020329

Eichler, K., Li, F., Litwin-Kumar, A., Park, Y., Andrade, I., Schneider-Mizell, C. M., … Gerber, B. (2017). The complete connectome of a learning and memory centre in an insect brain. Nature, 548(7666), 175–182. doi:10.1038/nature23455

Frank, D. D., Jouandet, G. C., Kearney, P. J., Macpherson, L. J., & Gallio, M. (2015). Temperature representation in the *Drosophila* brain. Nature, 519(7543), 358–361. doi:10.1038/nature14284

Gianluigi, M., Amit, D. J., & Nicolas, B. (2003). Retrospective and prospective persistent activity induced by Hebbian learning in a recurrent cortical network. European Journal of Neuroscience, 18(7), 2011–2024. doi:10.1046/j.1460-9568.2003.02908.x

Goldberg, J. H., Gabor, T., Dmitriy, A., & Rafael, Y. (2003). Calcium microdomains in aspiny dendrites. Neuron, 40(4), 807–821. doi:10.1016/S0896-6273(03)00714-1

Grimes, W. N., Zhang, J., Graydon, C. W., Kachar, B., & Diamond, J. S. (2010). Retinal parallel processors: More than 100 independent microcircuits operate within a single interneuron. Neuron, 65(6), 873–885. doi:10.1016/j.neuron.2010.02.028

Guo, A., Zhang, K., Ren, Q. Z., Su, H. F., & Chen, N. N. (2016). Functional connectivity mapping of decision-making in *Drosophila melanogaster*. In R. Wang & X. Pan (Eds.), Advances in Cognitive Neurodynamics *(V)* (pp. 35–40). Singapore: Springer Singapore.

Haga, T., & Fukai, T. (2018). Recurrent network model for learning goal-directed sequences through reverse replay. eLife, 7, e34171. doi:10.7554/eLife.34171

Hayworth, K. J., Kasthuri, N., Schalek, R., & Lichtman, J. W. (2006). Automating the collection of ultrathin serial sections for large volume TEM reconstructions. Microscopy & Microanalysis, 12(S02), 86–87. doi:10.1017/S1431927606066268

Heymann, J. A. W., Hayles, M., Gestmann, I., Giannuzzi, L. A., Lich, B., & Subramaniam, S. (2006). Site-specific 3D imaging of cells and tissues with a dual beam microscope. Journal of Structural Biology, 155(1), 63–73. doi:10.1016/j.jsb.2006.03.006

Inada, K., Tsuchimoto, Y., & Kazama, H. (2017). Origins of cell-type-specific olfactory processing in the *Drosophila* mushroom body circuit. Neuron, 95(2), 357–367.e354. doi:10.1016/j.neuron.2017.06.039

Joesch, M., Mankus, D., Yamagata, M., Shahbazi, A., Schalek, R., Suissa-Peleg, A., … Sanes, J. R. (2016). Reconstruction of genetically identified neurons imaged by serial-section electron microscopy. eLife, 5, e15015. doi:10.7554/eLife.15015

Kageyama, G. H., & Meyer, R. L. (1987). Dense HRP filling in pre-fixed brain tissue for light and electron microscopy. Journal of Histochemistry and Cytochemistry, 35(10), 1127–1136. doi:10.1177/35.10.3624853

Kasthuri, N., Hayworth, K. J., Berger, D. R., Schalek, R. L., Conchello, J. A., Knowles-Barley, S., … Lichtman, J. W. (2015). Saturated reconstruction of a volume of neocortex. Cell, 162(3), 648–661. doi:10.1016/j.cell.2015.06.054

Ke, Z., Guo, J. Z., Peng, Y., Wang, X., & Aike, G. (2007). Dopamine-mushroom body circuit regulates saliency-based decision-making in *Drosophila*. Science, 316(5833), 1901–1904. doi:10.1126/science.1137357

Kirkhart, C., & Scott, K. (2015). Gustatory learning and processing in the *Drosophila* mushroom bodies. The Journal of Neuroscience, 35(15), 5950–5958. doi:10.1523/JNEUROSCI.3930-14.2015

Knott, G., Marchman, H., Wall, D., & Lich, B. (2008). Serial section scanning electron microscopy of adult brain tissue using focused ion beam milling. Journal of Neuroscience, 28(12), 2959–2964. doi:10.1523/JNEUROSCI.3189-07.2008

Kohonen, T., & Oja, E. (1976). Fast adaptive formation of orthogonalizing filters and associative memory in recurrent networks of neuron-like elements. Biological Cybernetics, 21(2), 85–95. doi:10.1007/BF01259390

Larsen, C. W. (2003). Segment boundary formation in *Drosophila* embryos. Development, 130(23), 5625–5635. doi:10.1242/dev.00867

le Magueresse, C., Alfonso, J., Khodosevich, K., Martin, A. A., Bark, C., & Monyer, H. (2011). “Small axonless neurons”: Postnatally generated neocortical interneurons with delayed functional maturation. The Journal of Neuroscience, 31(46), 16731–16747. doi:10.1523/JNEUROSCI.4273-11.2011

Lee, T., Lee, A., & Luo, L. (1999). Development of the *Drosophila* mushroom bodies: Sequential generation of three distinct types of neurons from a neuroblast. Development, 126(18), 4065–4076.

Lei, Z., Chen, K., Li, H., Liu, H., & Guo, A. (2013). The GABA system regulates the sparse coding of odors in the mushroom bodies of *Drosophila*. Biochemical and Biophysical Research Communications, 436(1), 35–40. doi:10.1016/j.bbrc.2013.05.036

Li, H., Li, Y., Lei, Z., Wang, K., & Guo, A. (2013). Transformation of odor selectivity from projection neurons to single mushroom body neurons mapped with dual-color calcium imaging. Proceedings of the National Academy of Sciences of the United States of America, 110(29), 12084–12089. doi:10.1073/pnas.1305857110

Lin, A. C., Bygrave, A. M., de Calignon, A., Lee, T., & Miesenbock, G. (2014). Sparse, decorrelated odor coding in the mushroom body enhances learned odor discrimination. Nature Neuroscience, 17(4), 559–568. doi:10.1038/nn.3660

Liu, Q., Dang, C., & Cao, J. (2010). A novel recurrent neural network with one neuron and finite-time convergence for k-winners-take-all operation. IEEE Transactions on Neural Networks, 21(7), 1140–1148. doi:10.1109/TNN.2010.2050781

Liu, X., & Davis, R. L. (2009). The GABAergic anterior paired lateral neuron suppresses and is suppressed by olfactory learning. Nature Neuroscience, 12(1), 53–59. doi:10.1038/nn.2235

Macpherson, L. J., Zaharieva, E. E., Kearney, P. J., Alpert, M. H., Lin, T. Y., Turan, Z., … Gallio, M. (2015). Dynamic labelling of neural connections in multiple colours by trans-synaptic fluorescence complementation. Nature Communications, 6, 10024. doi:10.1038/ncomms10024

Mao, Z., & Davis, R. L. (2009). Eight different types of dopaminergic neurons innervate the *Drosophila* mushroom body neuropil: Anatomical and physiological heterogeneity. Frontiers in Neural Circuits, 3, 5. doi:10.3389/neuro.04.005.2009

Marta, R. A., Vitaladevuni, S. N., Yuriy, M., Yuriy, M., Zhiyuan, L., Shin-Ya, T., … de Polavieja, G. G. (2011). Wiring economy and volume exclusion determine neuronal placement in the *Drosophila* brain. Current Biology, 21(23), 2000–2005. doi:10.1016/j.cub.2011.10.022

Olsen, S. R., & Wilson, R. I. (2008). Lateral presynaptic inhibition mediates gain control in an olfactory circuit. Nature, 452(7190), 956–960. doi:10.1038/nature06864

Osinski, B. L., & Kay, L. M. (2016). Granule cell excitability regulates gamma and beta oscillations in a model of the olfactory bulb dendrodendritic microcircuit. Journal of Neurophysiology, 116(2), 522–539. doi:10.1152/jn.00988.2015

Papadopoulou, M., Cassenaer, S., Nowotny, T., & Laurent, G. (2011). Normalization for sparse encoding of odors by a wide-field interneuron. Science, 332(6030), 721–725. doi:10.1126/science.1201835

Pitman, J. L., Huetteroth, W., Burke, C. J., Krashes, M. J., Lai, S. L., Lee, T., & Waddell, S. (2011). A pair of inhibitory neurons are required to sustain labile memory in the *Drosophila* mushroom body. Current Biology, 21(10), 855–861. doi:10.1016/j.cub.2011.03.069

Rall, W. (1969). Time constants and electrotonic length of membrane cylinders and neurons. Biophysical Journal, 9(12), 1483–1508. doi:10.1016/S0006-3495(69)86467-2

Ren, Q., Li, H., Wu, Y., Ren, J., & Guo, A. (2012). A GABAergic inhibitory neural circuit regulates visual reversal learning in *Drosophila*. The Journal of Neuroscience, 32(34), 11524–11538. doi:10.1523/JNEUROSCI.0827-12.2012

Scholz-Kornehl, S., & Schwarzel, M. (2016). Circuit analysis of a *Drosophila* dopamine type 2 receptor that supports anesthesia-resistant memory. The Journal of Neuroscience, 36(30), 7936–7945. doi:10.1523/JNEUROSCI.4475-15.2016

Shu, X., Lev-Ram, V., Deerinck, T. J., Qi, Y., Ramko, E. B., Davidson, M. W., … Mcintosh, J. R. (2011). A genetically encoded tag for correlated light and electron microscopy of intact cells, tissues, and organisms. PLoS Biology, 9(4), e1001041. doi:10.1371/journal.pbio.1001041

Tanaka, N. K., Tanimoto, H., & Ito, K. (2008). Neuronal assemblies of the *Drosophila* mushroom body. The Journal of Comparative Neurology, 508(5), 711–755. doi:10.1002/cne.21692

Tomer, R., Denes, A. S., Tessmar-Raible, K., & Arendt, D. (2010). Profiling by image registration reveals common origin of annelid mushroom bodies and vertebrate pallium. Cell, 142(5), 800–809. doi:10.1016/j.cell.2010.07.043

Wang, X. J. (2008). Decision making in recurrent neuronal circuits. Neuron, 60(2), 215–234. doi:10.1016/j.neuron.2008.09.034

Wilson, R. I., Turner, G. C., & Laurent, G. (2004). Transformation of olfactory representations in the *Drosophila* antennal lobe. Science, 303(5656), 366–370. doi:10.1126/science.1090782

Wong, K.-F. (2006). A recurrent network mechanism of time integration in perceptual decisions. Journal of Neuroscience, 26(4), 1314–1328. doi:10.1523/JNEUROSCI.3733-05.2006

Wu, C. L., Shih, M. F., Lai, J. S., Yang, H. T., Turner, G. C., Chen, L., & Chiang, A. S. (2011). Heterotypic gap junctions between two neurons in the *Drosophila* brain are critical for memory. Current Biology, 21(10), 848–854. doi:10.1016/j.cub.2011.02.041

Wu, J. S., & Luo, L. (2006). A protocol for dissecting *Drosophila melanogaster* brains for live imaging or immunostaining. Nature Protocols, 1(4), 2110–2115. doi:10.1038/nprot.2006.336

Wu, Y., Ren, Q., Li, H., & Guo, A. (2012). The GABAergic anterior paired lateral neurons facilitate olfactory reversal learning in *Drosophila*. Learning & Memory, 19(10), 478–486. doi:10.1101/lm.025726.112

Yang, C. H., Shih, M. F., Chang, C. C., Chiang, M. H., Shih, H. W., Tsai, Y. L., … Wu, C. L. (2016). Additive expression of consolidated memory through *Drosophila* mushroom body subsets. PLoS Genetics, 12(5), e1006061. doi:10.1371/journal.pgen.1006061

Yasuyama, K., Meinertzhagen, I. A., & Schurmann, F. W. (2002). Synaptic organization of the mushroom body calyx in *Drosophila melanogaster*. The Journal of Comparative Neurology, 445(3), 211–226. doi:10.1002/cne.10155

Zheng, Z., Lauritzen, J. S., Perlman, E., Robinson, C. G., Nichols, M., Milkie, D., … Sharifi, N. (2018). A complete electron microscopy volume of the brain of adult *Drosophila melanogaster*. Cell, 174(3), 730–743.e722. doi:10.1016/j.cell.2018.06.019

