## Supplemental for "The reconstruction and functional mapping of a recurrent microcircuit in *Drosophila* mushroom body"

### Supplemental methods

#### *Establishment of the EM data analysis procedure*

After obtaining continuous EM images of mushroom bodies, we needed to find APL neurons and their connections. This work involved collaborative processing of huge amounts of data (> 70GB) collected by multiple people, requiring an effective workflow.

As the basis of the workflow, we first determined the working principle of the multiperson collaboration and selected the appropriate software tools. The core consideration of using a multiperson collaborative method to analyze EM data is the reliability of the analysis results. An artificial recognition process inevitably produces errors, which may come from the uncertainty of the image data or human preferences. To minimize the impact of human preferences on the analysis results, a multiperson collaborative proofreading and checking process is essential. We compared several kinds of software (Amira, Reconstruct (Fiala, 2005), TrakEM2 (Cardona et al., 2010)) that can be used for EM reconstruction and chose to use TrakEM2. This program is based on a widely used open source image processing program Fiji (Schindelin et al., 2012) based on ImageJ (Schneider, Rasband, & Eliceiri, 2012), which has rich support documents to quickly guide our solutions to problems during its use; second, TrakEM2 provides a program interface that conveniently processes reconstructed data by itself, facilitating the expansion and flexibility of its use. Finally, there are also many support documents for this software. Due to the open source nature of the program, procedural professionals can help you overcome possible problems and expand the functions of the software. In summary, TrakEM2 software provides a suitable tool for small-scale (approximately 5 people) neural metadata reconstruction because of its convenient use, easy expansion and easy-to-find support features.

The first step in the workflow was the alignment of consecutive slice images (or alignment). Neuron tracking requires accurate alignment of successive upper and lower images. For technical reasons, slight displacement is inevitable with successive FIB-SEM image acquisition, which requires efficient methods to align these images one by one. With the help of a mature image alignment algorithm, we obtained regularly aligned consecutive EM images. The aligned images revealed complete and continuous cell morphology on all sections of the three-dimensional space (Fig. S1).

The second step in the workflow was neuron reconstruction. In this step, we only recognized and reconstructed the target APL neuron and the neurons that formed synapses with it, thereby adopting the sparse reconstruction strategy. This strategy avoids reconstructing all neurons in the vertical leaves of the mushroom body, thereby limiting the workload tremendously as there are approximately 1300 (Aso et al., 2009) KCs in vertical leaves alone. If no other means such as labeling were used, we would need to reconstruct all neurons to inspect and judge the morphological characteristics of cells and identify the target neurons; even so, there would still be a risk of misjudgment, because we do not know the fine morphological structure of APL neurons, and there may be other unknown morphological similarities or neuronal interference. The above problem can be avoided by using an HRP labeling strategy for membrane localization, wherein the deepening of the membrane color in EM images is a direct indication of the location of APL neurons. Therefore, in our workflow, APL neurons were identified by using cross-sections of membrane-deepened neurons as seeds. Through synaptic recognition, APL connective neurons were found, and the reconstruction of APL local connections was completed (Fig. S2).

In the reconstruction process, we first identified APL neurons with membrane markers (Fig. S2). We identified and searched the cell membranes deepened in the data set (Fig. 1A) and tracked the identified neuron cross-sections in the Z direction. We found that HRP-labeled neurons had obvious membrane deepening on some sections (Fig. 1B) but not on other sections, indicating that HRP was not uniformly distributed throughout the cell membrane. Therefore, we also established criteria for identifying APL neurons: (Fiala, 2005) obvious membrane deepening on a single EM image (Cardona et al., 2010) or traced neurons with membrane deepening on certain sections. In the second step, we tracked the direction of neurons in continuous images and labeled them in the software to obtain the orientation skeleton of the APL neuron (Fig. S3B). The third step was to identify the synapses of neurons. Using a similar recognition-proofreading process for reconstructing neurons, we identified neuronal synapses. The synaptic structure of *Drosophila melanogaster* is easily recognized in EM images, and no special marking is required. There are three elements for synaptic recognition in EM images: a presynaptic T-bar; synaptic vesicles; and postsynaptic membrane deepening. Because the membrane labeling method we used interferes with the deepening of postsynaptic membranes, in this data set, to reduce class I errors (false positives), we only identified synapses using the T-bar, likely underestimating the number of synapses. In the fourth step, we reconstructed the volume of neurons. By identifying the boundaries of neurons on each section and painting the area occupied by them, we finally obtained the detailed morphology of neurons in three-dimensional space (Fig. S3C). Among these steps, the fourth step of painting is the most time-consuming; we examined some automatic recognition tools; however, in this data set, their recognition ability was not sufficient, and they were unable to reduce the amount of manual labor needed; therefore, we performed this step using the digital drawing board as well as human labor .

After tracing and reconstructing APL neurons, we carried out proofreading and error correction using a multiperson collaboration. The core problem of proofreading is the lack of a completely correct "truth value", and each recognizer may make different judgments. As a means of approaching the true value, the common method is to identify the same part by many people and then to examine whether an agreement can be reached; if a certain recognition result cannot reach agreement, it is discarded. In this way, the authenticity of the checked results was ensured as much as possible; at the same time, there may be a few omissions in the results. This underestimating proofreading process is actually a trade-off between uncertainty, which can be analogous to reducing class I errors (false positive, identifying nonexistent neurons) and inevitably increasing class II errors (false negative, neurons are not recognized). In the subsequent data analysis and interpretation, we believe that class I errors are far more misleading than class II errors; therefore, we tried to avoid class I errors in proofreading. Specifically, as shown in Fig. S2B, each person in the working group first checked the identified data and identified possible errors and omissions in a part of the data. They then integrated the information and redistributed it to each person for voting on the basis of the aforementioned criteria to decide whether to accept the modification. The above process was then repeated for the updated data until no more errors or omissions were found. To meet such process requirements, we cooperated with software experts to modify the software and develop program scripts to assist with the accurate execution of the process. The whole proofreading process covered each identified neuron and synapse at least three times, and the coverage of many areas reached more than five iterations. Through this recognition-proofreading process, the most accurate reconstructed data of EM-visualized neurons was obtained.

In combination with the above methods, we organized a multiperson collaboration group with 4 to 6 people for simultaneous data reconstruction. First, APL neurons in the collected data from the vertical lobe of the mushroom body were identified and reconstructed. Then, their output synapses (postsynapses) and input synapses (presynapses) were identified. Then, some output and input APL neurons were tracked through these synapses. All of these data provided more details on their morphology and a solid foundation for their connection characteristics.

Notably, we attempted the automatic recognition method for neuron reconstruction. We compared and evaluated the effects of artificial recognition and automatic recognition (Hui Li, *unpublished*) based on a machine learning algorithm. On the surface, manual recognition is time-consuming and laborious, requiring repetitive work, whereas computer algorithms appear efficient and convenient. However, the computer algorithm results in some false-positive recognition, that is, a portion of an image belonging to a different neuron is classified as belonging to the same neuron. Despite the efforts of computer algorithm experts working with us to improve their accuracy and reduce the occurrence of false positives, some mistakes are inevitable, partly because the computer algorithm can only process the whole image, and the workload of recognition is enormous. To check the recognition results of the whole image, many human resources are needed, even more than are needed with manual recognition from the beginning. At present, the most advanced computer image recognition algorithm can only process specially simplified neurons (Moritz et al., 2013), far less than the image recognition ability of the brain. Therefore, we abandoned the use of automated neuron recognition methods and adopted another strategy: through collaboration with computer software engineering experts, we can improve the operation efficiency of software tools, reduce the simple and repetitive workload, and enable users of software tools to focus more on complex neuron recognition problems.

***Synapse arrangement index, AI***

The synapses on the neurites were grouped according to their directions. The grouping of a continuous input synapse into an input cluster and a continuous output synapse into an output cluster results in many input clusters and output clusters. For a direction D (input or output), there are a total of N*d* synapses distributed in G*d* clusters. Then, the synaptic arrangement index AI*d* in this direction was calculated by the following formula:

$$\mathrm{AI}d = \begin{cases} \dfrac{Gd - 1}{Nd - 1}, & Nd > 1 \\ 1, Nd = 1 \end{cases}$$

In this way, the AIin of input synapse and the AIout of output synapse were obtained. In the data analysis of APL input and output neurons, neurites of N*d* < 3 were not displayed (Fig. S7).

***Statistical analysis of synapse number under light microscopy***

Data after deconvolution processing were imported into Fiji software, and synaptic counts were performed using modified dissector TrakEM2, which could mark successive layers of staining. We used synaptobrevin-GFP to label presynaptic vesicles. In the image, synapses were identified by staining spanning several layers of the image at the same time, and noise was identified by discontinuous staining. Based on the above criteria, we used automatic computer identification and manual identification to count the number of synapses.

**Results**

#### *APL EM reconstruction results*

APL neurons were reconstructed in the vertical lobe of the mushroom body (Table 1) using the previously established continuous EM method and data analysis process with labeled cells. We observed that the APL neurons penetrated the inside of the vertical lobe of the mushroom body and dispersed more evenly in the horizontal direction, and the top of the APL neuron was more sparse in the vertical direction than in the middle (Fig. S4). A large proportion of these neurites extended in the vertical direction (Fig. S5), which is consistent with the direction of the mushroom body neurons around them. This directionality can be inferred from the cross-section of the neurites in most of the EM sections. Near the top of the image set, most of these neurites are bent and inclined to the horizontal direction. Notably, there are less than 20 APL branches in the horizontal section of the α lobe (Fig. S4), and approximately 1000 (Aso et al., 2009) KCs in the α lobe; less than 30 APL branches in the α lobe (Fig. S4), and approximately 300 (Aso et al., 2009) KCs in the α lobe. Therefore, the average number of APL branches in the α lobe needs to be at least 50 to cover all KCs, while the number of branches in the α lobe needs at least 10. In our reconstructed results, the total number of input synapses in the α lobe was only approximately 100 (Table 1), which is far less than the total number of input synapses needed to cover all KCs; even considering that our recognition results underestimate the true situation by four to five times (Fig. S6), there are not enough input synapses to cover all KCs. Similar conditions exist in the output synapses of APL neurons in the α lobe and in the α’ lobe. Thus, in the vertical lobe, APL neurons do not seem to act as broad inhibitory neurons, broadcasting inhibitory signals to all KCs, unlike the similar neuron found in locusts, the GGN (Papadopoulou, Cassenaer, Nowotny, & Laurent, 2011), suggesting that we may not directly apply the physiological characteristics of GGNs to APL neurons.

***Presynapses and postsynapses on APL neurites are intermixed***

Synapses are important places for information transmission, and their distribution can indicate the information flow of neurons. The close alignment of input and output synapses indicates that neurons process information locally (e.g., A17 neurons (Grimes, Zhang, Graydon, Kachar, & Diamond, 2010)), which is an important indicator of neuron function. APL reconstruction and synaptic connection analysis revealed that input synapses and output synapses are intermixed on the neurite (Fig. S7); with the extension of the neurite, there are often several continuous input synapses followed by several output synapses. The location of the input and output synapses also coincides with the alternating distribution of enlarged and thin sections of the neurite mentioned above. To better describe this phenomenon, synapses were divided into different clusters, each of which with only one continuous synapse in the same direction (Fig. S7). Thus, the "input cluster" consisting of input synapses and the "output cluster" consisting of output synapses were alternately arrange on neurons. At the same time, to investigate the mixed arrangement of synapses in different directions and compare across different neurons, the distance between synapses and the length and branches of neurons had to be disregarded, and the topological arrangement between them had to be extracted. Here, we used an "arrangement index" (AI) to achieve this goal. We clustered synapses in a certain direction according to the above method, compared the total number of synapses with the number of clusters, and normalized them between 0 and 1, providing a degree of the convergence/dispersion of synapses. For example, when the input synapse and the output synapse are arranged alternately at 1:1 intervals, the AIin of the input synapse and the AIout of the output synapse are both 1, with the highest degree of mixing; correspondingly, for example, in ideal mammalian bipolar neurons, the input synapses are concentrated in one region while the output synapses are concentrated in another region, resulting in an AIin and AIout of 0, and the mixing degree is the lowest. Based on APL synaptic data in the vertical lobe, we obtained the AIin and AIout of APL neurons, showing the extent of the arrangement of input and output synapses (Fig. S7C). Compared with A17 neurons in the mammalian retina (Grimes et al., 2010), the synaptic alignment of APL synapses is slightly less mixed (shown in yellow in Fig. S7C).

### References


Aso, Y., Grubel, K., Busch, S., Friedrich, A., Siwanowicz, I., & Tanimoto, H. (2009). The mushroom body of adult *Drosophila* characterized by GAL4 drivers. *Journal of Neurogenetics, 23*, 156-172. doi:10.1080/01677060802471718

Cardona, A., Saalfeld, S., Preibisch, S., Schmid, B., Cheng, A., Pulokas, J., . . . Hartenstein, V. (2010). An integrated micro- and macroarchitectural analysis of the *Drosophila* brain by computer-assisted serial section electron microscopy. *PLoS Biology, 8*(10), e1000502. doi:10.1371/journal.pbio.1000502

Fiala, J. C. (2005). Reconstruct: A free editor for serial section microscopy. *Journal of Microscopy, 218*(1), 52-61. doi:10.1111/j.1365-2818.2005.01466.x

Grimes, W. N., Zhang, J., Graydon, C. W., Kachar, B., & Diamond, J. S. (2010). Retinal parallel processors: More than 100 independent microcircuits operate within a single interneuron. *Neuron, 65*(6), 873-885. doi:10.1016/j.neuron.2010.02.028

Moritz, H., Briggman, K. L., Turaga, S. C., Viren, J., Sebastian, S. H., & Winfried, D. (2013). Connectomic reconstruction of the inner plexiform layer in the mouse retina. *Nature, 500*(7461), 168. doi:10.1038/nature12346

Papadopoulou, M., Cassenaer, S., Nowotny, T., & Laurent, G. (2011). Normalization for sparse encoding of odors by a wide-field interneuron. *Science, 332*(6030), 721-725. doi:10.1126/science.1201835

Schindelin, J., Argandacarreras, I., Frise, E., Kaynig, V., Longair, M., Pietzsch, T., . . . Schmid, B. (2012). Fiji: An open-source platform for biological-image analysis. *Nature Methods, 9*(7), 676-682. doi:10.1038/nmeth.2019

Schneider, C. A., Rasband, W. S., & Eliceiri, K. W. (2012). NIH Image to ImageJ: 25 years of image analysis. *Nature Methods, 9*(7), 671-675. doi:10.1038/nmeth.2089

### Supplementary Figures

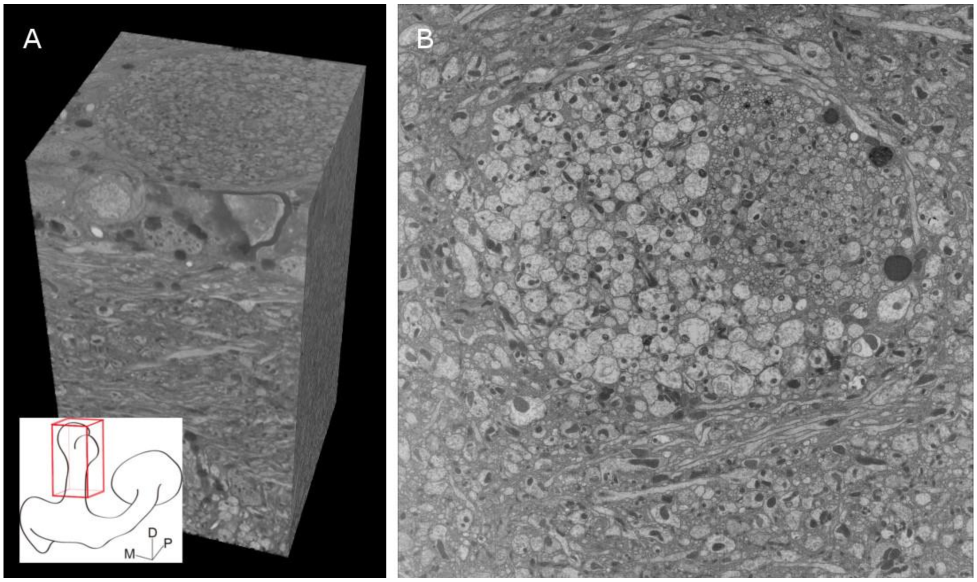


**Fig. S1 The vertical lobe mushroom body dataset and a section of the α and α′ lobes**

**(A)** 3D view of a portion of the continuous EM data after alignment showing the vertical lobe of the mushroom body as indicated in the left bottom red cube. There are a total of 1300 EM images in the left EM dataset, occupying an approximately 33(x)×37(y)×52(z) $\mu m^3$ space. Notably, the side of the cube in the picture is not an image that was collected directly but the result of alignment and stitching. The side image, similar to direct slicing, shows that the acquisition and alignment of the image set was very successful. **(B)** A slice in the data set is shown and is approximately 35 µm away from the top of the vertical lobe, occupying a 30×30 $\mu m^2$ area horizontally. The densely arranged vertical lobe neurons of the mushroom body can be seen in the image, and the obvious larger cells are cross sections of α' neurons. The resolution in the XY direction and in the Z direction is 7 nanometers and 40 nanometers, respectively.

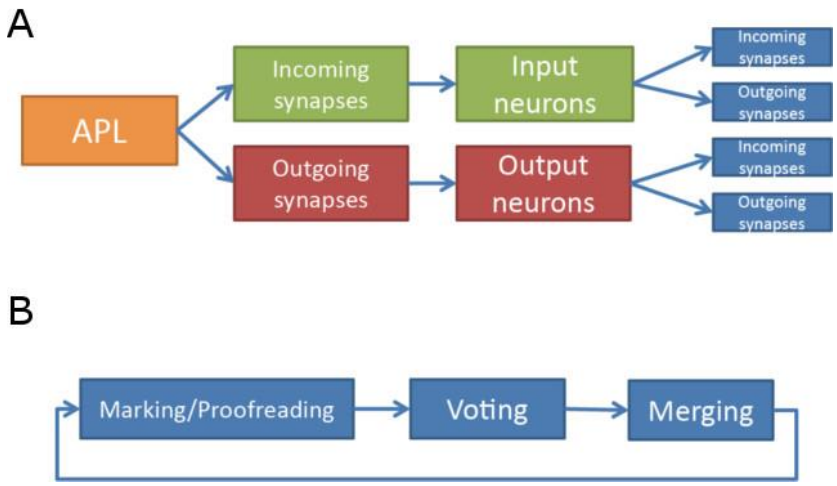


**Fig. S2 Workflow of the reconstruction of an APL neuron and its connectomes**

**(A)** The strategy for reconstructing APL neurons is demonstrated. First, the APL neurons is recognized and reconstructed. Then, the input and output synapses of the APL neuron are identified. These synapses correspond to the cross sections of input and output neurons. Then, their directions are traced from these cross sections. Finally, the synapses of input and output neurons are identified. This is a sparse reconstruction process that traces the first cascade of APL neurons to reconstruct and analyze the data more quickly. **(B)** The proofreading strategy for reconstruction, performed at every step. Specifically, synapses or neurons are identified or tracked first, and existing data are checked and proofread. The results of these proofreading steps are then distributed to different proofreaders for voting. The voting results are integrated, and the existing reconstructed data are updated. The process is then executed in a loop. After several rounds of proofreading, new errors will no longer be found, and the proofreading process will be complete.

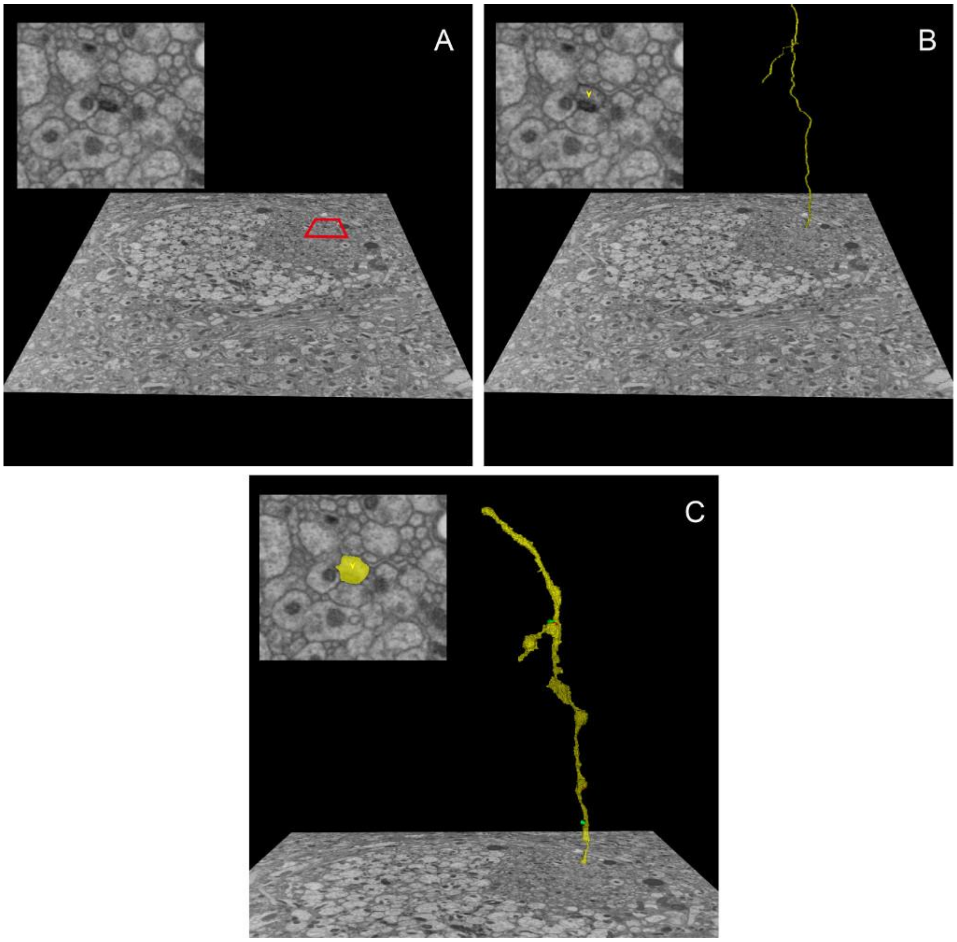


**Fig. S3 APL neuron 3D reconstruction procedure**

This is a typical process of neuronal remodeling of an APL neuron.

**(A)** First, the cross sections of neurons in single-layer images are identified as a node, and the neurons with deeper membranes are identified as APL neurons. The top left image is an enlargement of the picture in the red box. **(B)** The direction of neurons is then tracked. The yellow skeleton shows the artificial marking of APL neurites tracked in 3D space. The skeleton is composed of many artificially labeled nodes, which are labeled as a part of a neuron skeleton of on each layer of EM data. A cross-section image of an APL neurons generally has one node. The upper left image is an enlargement of the picture in the red box in (A) in (B). The small yellow arrow near the center of the figure indicates the nodes marked on the section of the layer.

**(C)** The cross-sectional area of each image layer occupied by neurons (e.g., yellow in the upper left picture) gives the volume in three dimensions (yellow in the larger picture). The input and output synapses (green and red cones in the larger picture) of the APL neuron can be obtained by identifying synaptic structures on neurons. Synaptic recognition can also be performed at the same time as skeleton tracking (B). The final result is the morphological remodeling of the APL neurites and their synaptic connections.

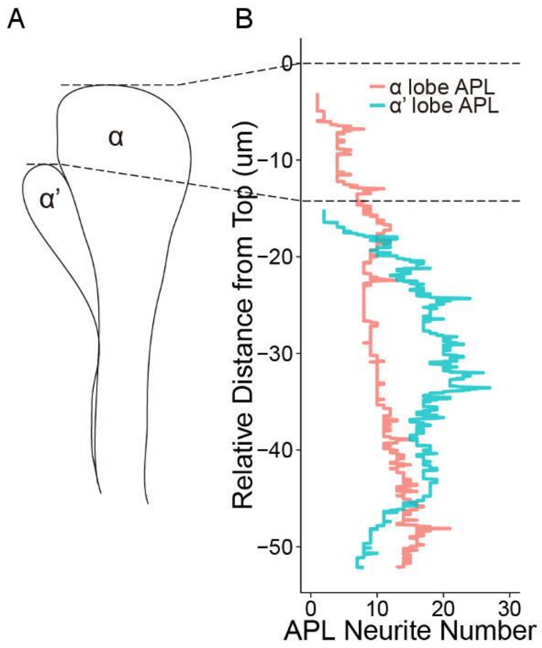


**Fig. S4 Distribution of APL neurons in the vertical lobe of the mushroom body**

The branching of APL neurons shown by EM reconstruction. The number of APL sections of each layer in a continuous image is indicated by **(B)** figure. At the bottom of the data set, i.e., the ventral side of the vertical lobe of the mushroom body, the number of APL neurons in the sections of the α lobe and α′ lobe was approximately 20; in the middle of the vertical lobe, the number of sections in the α lobe was more than that in the α lobe; at the top of the vertical lobe, the distribution of APL neurites in the α lobe and α′ lobe is obviously less. The number of APL neurites in the figure is represented by the number of nodes in the reconstructed skeleton and therefore sometimes contains different nodes of the same lateral branch at the same horizontal level, indicated by the spikes in the figure.

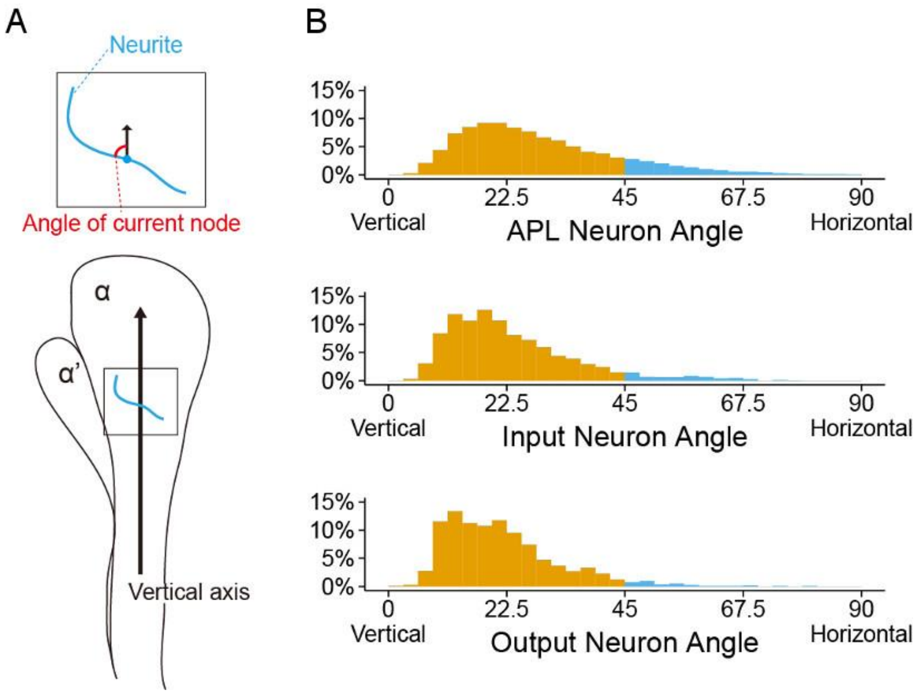


**Fig. S5 Trends of APL neurons and their input-output neurons in the α lobe**

The orientation of reconstructed α intralobar neurons.

**(A)** A schematic view of the node angle on the neurite process. The neurites (shown in sky blue) walk in the α lobe and form an angle with the vertical axis (shown in red). The angle distribution (B) of all reconstructed neurons was obtained by statistical analysis. **(B)** The angle distribution of the neurites in the upper and middle parts of the APL neurons shows that most of the neurites are vertical (yellow) and some are horizontal (blue). (B) The middle and lower parts of the graph are the orientation of input neurons and output neurons to the APL neuron, respectively. Obviously, the direction of input and output neurons is more vertical than that of the APL neuron, and the peak of the angle distribution curve is steeper and more towards 0 degrees (i.e., the vertical direction).

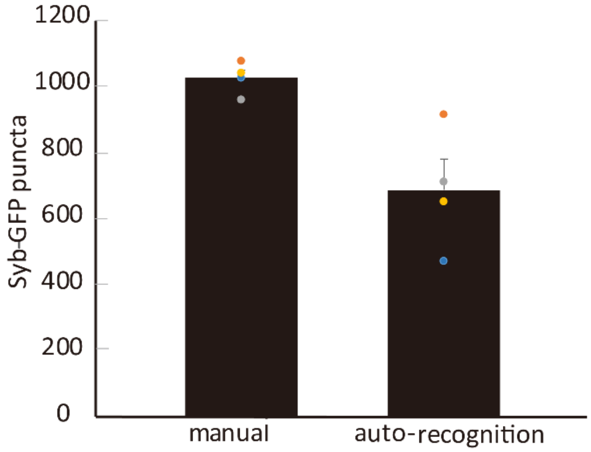


**Fig. S6 Approximately 600~1000 APL presynapses labeled by synaptobrevin-GFP under light microscopy.** The APL presynapses are labeled by synaptobrevin-GFP. The points with the same color represent the APL data of the same fly. There are four flies in total shown here with data from the vertical lobe.

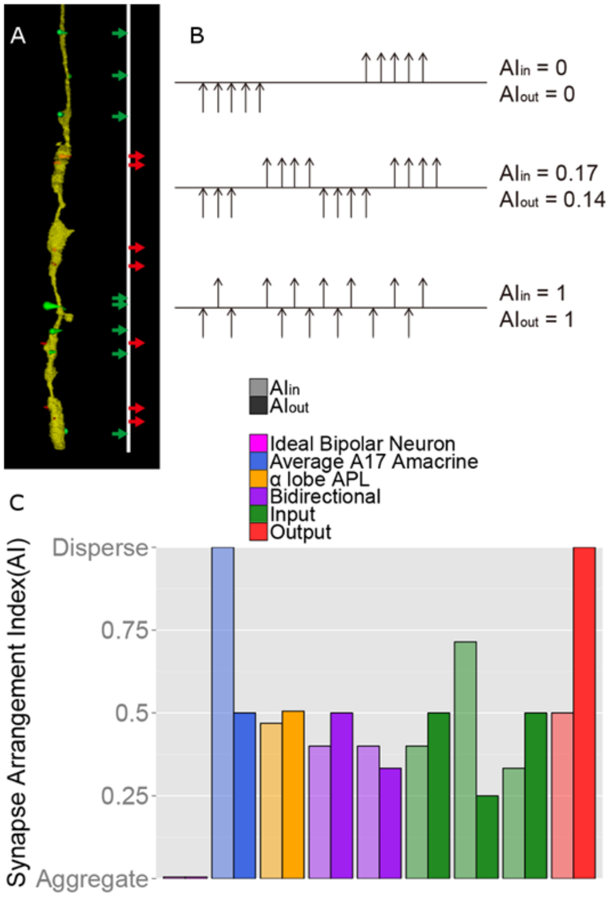

**Fig. S7 Synapse arrangement of APL neurons and amacrine A17 neurons**

**(A)** The reconstruction data of APL neurites in the α lobe. The reconstruction data of APL neurites in the α lobe show the typical synaptic mixing arrangement. Yellow is the shape of the neurite, the small green cone attached to it is the input synapse, and the small red cone is the output synapse. To the right of the neuron is an image that shows the synaptic distribution on the neurite, and the mixed arrangement of the input and output synapses is easy to see. **(B)** The AI values under different synaptic arrangements are indicated. The upper part shows that the input and output synapses are clustered into two separate groups, such as typical bipolar cells, where AIin and AIout are both 0; the lower part shows that the input and output synapses are alternately arranged one by one, when AIin and AIout are both 1; and the middle part shows a mixed arrangement of synapses in both directions. In this case, AIin is 0.17 and AIout is 0.14. **(C)** The synaptic AI of different neurons. The AIin and AIout were calculated for each identified neuron and are represented by a pair of cylinders with the same color but different saturation. AI was calculated by ignoring the distance factor and retaining only the information of the topological structure for comparison among different neurons. The data of an ideal bipolar neuron assume that the input and output synapses are separated into two independent groups, making AIin and AIout both 0. On average, there is one input synapse to every two output synapses in an A17 amacrine cell, and the whole is composed of many such units (Grimes et al., 2010). Based on this, AIin is estimated to be 1 and AIout to be 0.5. Yellow indicates the APL neurites in the α lobe, purple indicates the input and output neurons in the α lobe, green indicates the input neurons in the α lobe, and red indicates the output neurons in the α lobe. The data here do not include neurons with fewer than three synapses in either direction.

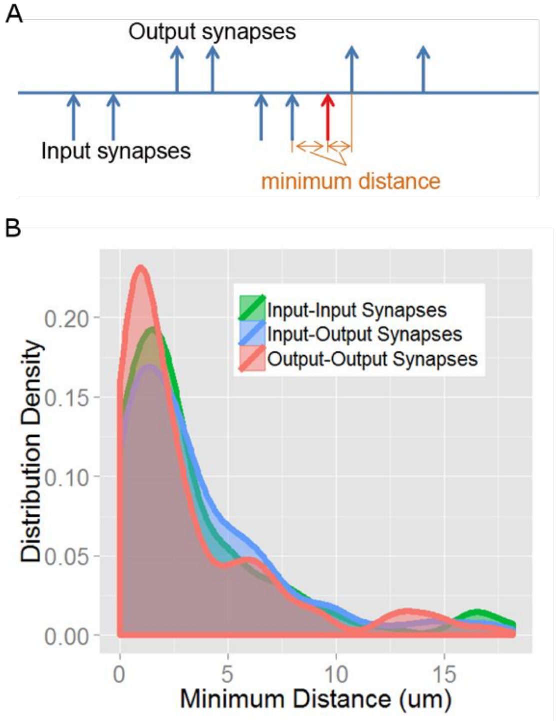


**Fig. S8 Minimum distance between input and output synapses of APL neurons**

**(A)** The minimum distance between synapses is the distance between a synapse and its nearest synapse. The synapse shown in red is an input synapse. Its nearest input synapse is on the left. The distance between them is the distance between input-input synapses. The distance between the nearest output synapses and the red synapse is on the right side. The distance between them is classified as the distance of input-output synapses. A similar categorization was extended to output-output synapses. **(B)** Distribution of the minimum distance of APL synapses of the different groups in the α lobe. The red curve shows the distribution of the minimum distance between output synapses. The peak value of the distribution curve is approximately 2 μm and extends to 15 μm, which indicates that a large number of synapses are very close. The green curve indicates the minimum distance of input-input synapses and is similar to the distribution of output-output synapses. Notably, the distribution of the blue curve indicating the minimum distance between input and output synapses is also similar to those of the first two groups, indicating that the distance between input and output synapses is approximately equal to the distance between input and output synapses. These data also confirm the mixed arrangement pattern of input and output synapses in APL neurons.

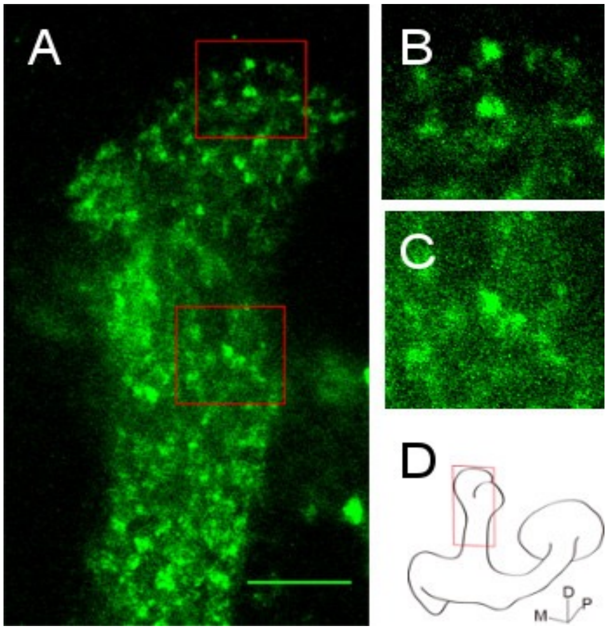


**Fig. S9 Light imaging showing that APL presynapses are located on the enlarged part of the neurite in the vertical lobe of *GH146-Gal4/+; UAS-n-synaptobrevin-GFP/+* flies**

**(A)** Confocal image of the vertical lobe of *GH146/+; UAS-n-syb-GFP/+* flies, displaying a specific fluorescence signal at presynapses of a frontal section (D). **(B)(C)** Enlarged images of the two squares in (A) from top to bottom, showing multiple fluorescent puncta. The scale bar is 10 μm. **(D)** Sketch of the frontal section of the left hemisphere of the mushroom body.

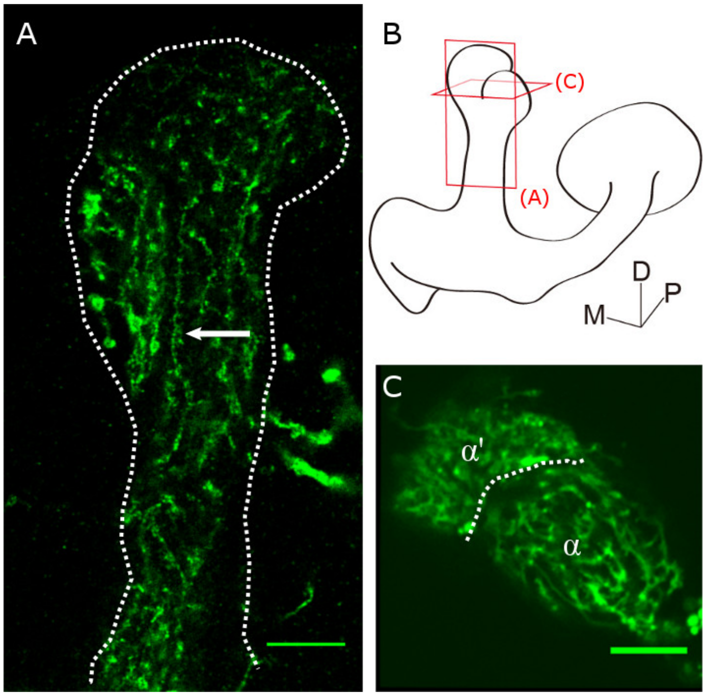


**Fig. S10 APL neurite showing the intermixed enlarged and thin regions in *GH146-Gal4/+; UAS-mGFP/+* flies**

**(A)** Confocal images of branches of APL neurites in a frontal section of the vertical lobe of the mushroom body in a *GH146-GAL4; UAS-mGFP* fly, as indicated in (B) (labeled with red A). Here, the white arrow displays the neurites along the vertical lobe. The olfactory PN is labeled outside of the dotted line. **(B)** Sketch of a frontal section (A) and horizontal section (C) of the left hemisphere of the mushroom body. **(C)** Confocal image of the horizontal section of APL neurites. The dotted line separates the α and α′ lobe. APL neurites are more concentrated in the α′ lobe than in the α lobe. The scale bar is 10 μm.

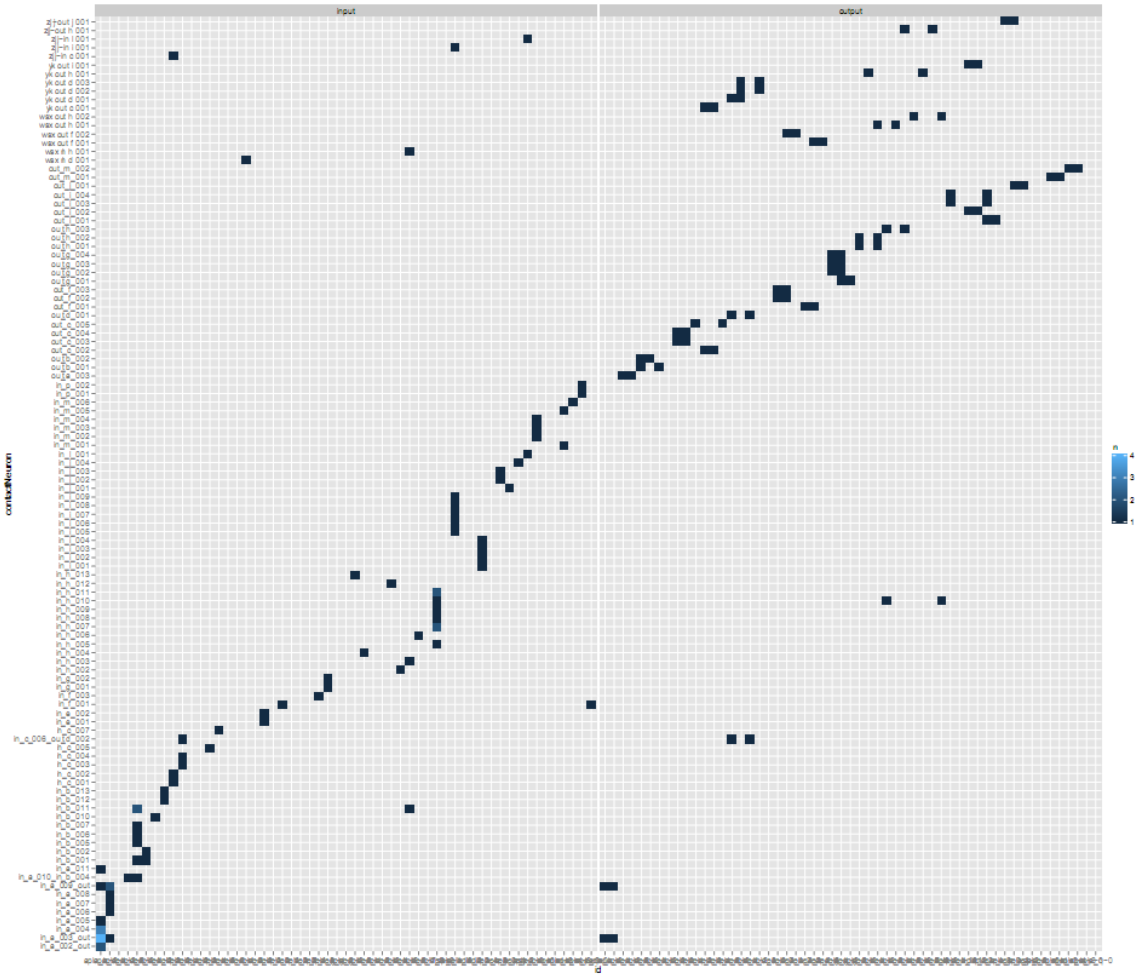


**Fig. S11 The connectome module between APL neurons and KCs is nonrepetitive and nonredundant**

The vertical axis displays all of the neurons connected with APL neurons in the vertical lobe. The intersection position indicates how many connections are between them, showing one-to-one connection relationship. The blue bar from deep to light indicates the number of connection from 1-4. This data indicates that more than 95% of the connections between APL neurons and its connecting neurons were nonrepetitive.

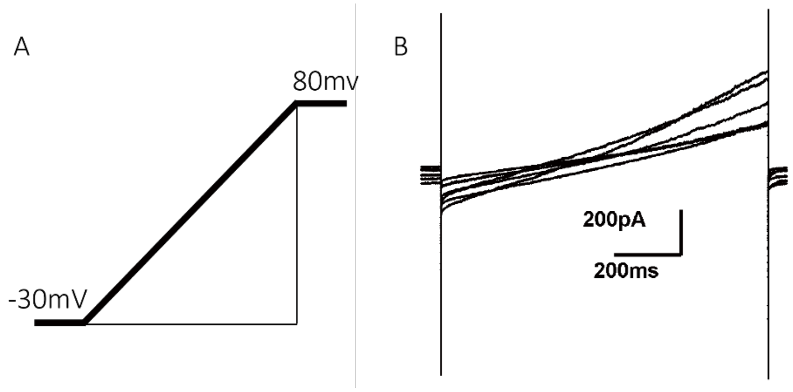


**Fig. S12 APL neuronal cell bodies elicit no action potential**

**(A)** The cell bodies of APL neurons were depolarized with stimulations ranging from -30 mV to 80 mV using whole-cell patch clamp recordings. **(B)** When APL neuronal cell bodies were depolarized, no action potentials appeared, as determined by the cell body currents.

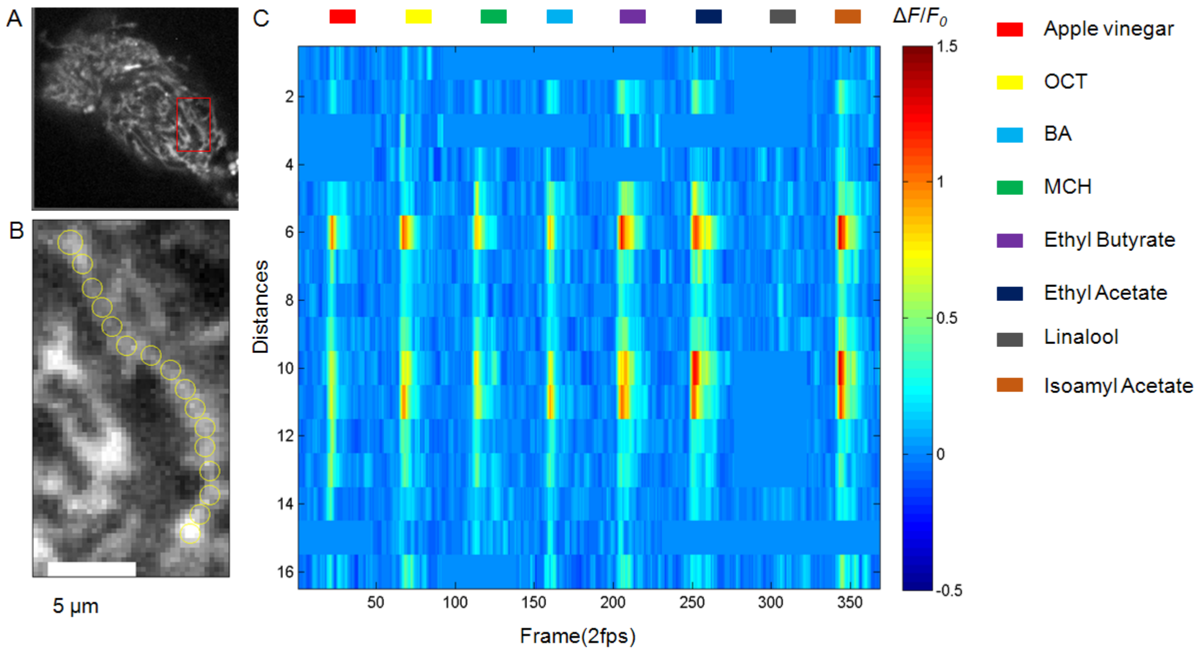


**Fig. S13 Example of the response of a single APL neurite calcium domain to eight different odors**

Odor stimuli were the same in all odor-delivery calcium imaging experiments

**(A)** Horizontal section of showing an APL neurite. **(B)** Magnification of a single APL neurite ROI shown in the red rectangle found in (A). **(C)** Heatmap of the single APL neurite (B) calcium imaging response to eight different odors. Vertical axis represents the neurite from bottom to top; each ROI is 2 μm, and the horizontal axis represents the scanning frame with rate of 2 fps.

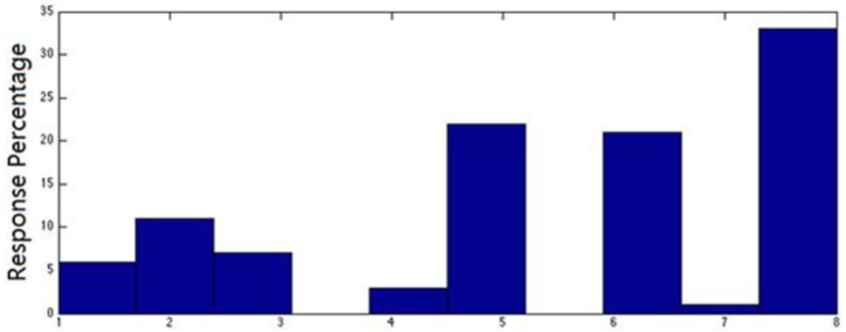


**Fig. S14 Different APL neurites have different preferences to eight odors**

Overall preferences to different odors for different APL neurites.

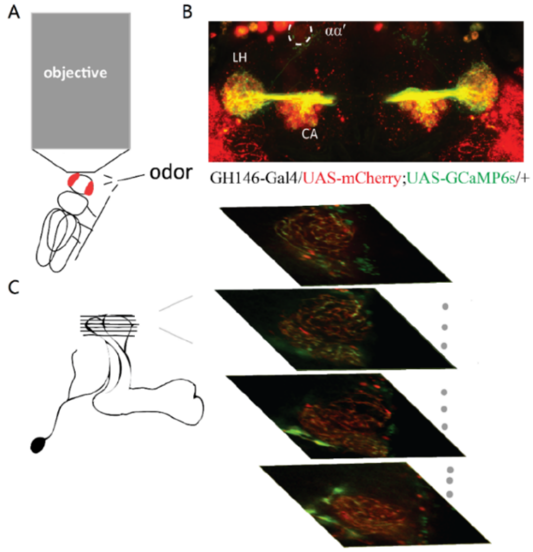


**Fig. S15 Experimental setup for calcium imaging in which flies were exposed to odor and electrical stimuli**

**(A)** Illustration of the imaging setup for odor and electrical stimuli. The fly stands upright on the electric copper plate during the experiment. **(B)** A scan view of a mushroom body and APL neurite. mCherry (in red) and GCaMP6s (in green) expression are driven by *GH146-Gal4*. *GH146-Gal4/UAS-mCherry; UAS-GCaMP6s/+*. αα′ represents the tip of the vertical lobe, CA represents the mushroom body calyx, and LH represents the lateral horn. **(C)** Imaging illustration of a multilayer scan of the vertical lobe, dual color APL neurites.
